# Faster-haplodiploid evolution under divergence-with-gene-flow: simulations and empirical data from pine-feeding hymenopterans

**DOI:** 10.1101/2021.04.09.439183

**Authors:** Emily E. Bendall, Robin K. Bagley, Vitor C. Sousa, Catherine R. Linnen

## Abstract

Although haplodiploidy is widespread in nature, the evolutionary consequences of this sex determination mechanism are not well characterized. Here, we examine how genome-wide hemizygosity and a lack of recombination in haploid males affects genomic differentiation in populations that diverge via natural selection while experiencing gene flow. First, we simulated diploid and haplodiploid “genomes” (500-kb loci) evolving under an isolation-with-migration model with mutation, drift, selection, migration, and recombination; and examined differentiation at neutral sites both tightly and loosely linked to a divergently selected site. So long as there is divergent selection and migration, sex-limited hemizygosity and recombination cause elevated differentiation (i.e., produce a “faster-haplodiploid effect”) in haplodiploid populations relative to otherwise equivalent diploid populations, for both recessive and codominant mutations. Second, we used genome-wide SNP data to model divergence history and describe patterns of genomic differentiation between sympatric populations of *Neodiprion lecontei* and *N. pinetum*, a pair of pine sawfly species (order: Hymenoptera; family: Diprionidae) that are specialized on different pine hosts. These analyses support a history of continuous gene exchange throughout divergence and reveal a pattern of heterogeneous genomic differentiation that is consistent with divergent selection on many unlinked loci. Third, using simulations of haplodiploid and diploid populations evolving according to the estimated divergence history of *N. lecontei* and *N. pinetum*, we found that divergent selection would lead to higher differentiation in haplodiploids. Based on these results, we hypothesize that haplodiploids undergo divergence-with-gene-flow and sympatric speciation more readily than diploids.

## Introduction

In terms of both species richness and biomass, haplodiploid organisms account for a substantial proportion of terrestrial biodiversity (Forbes et al., 2018; Hölldobler & Wilson, 1990). Haplodiploidy (arrhenotoky), a sex determination mechanism in which females develop from fertilized eggs and are diploid, while males develop from unfertilized eggs and are haploid, has evolved repeatedly in diverse arthropod lineages and is the mode of sex determination for an estimated 12% of extant animal species (Blackmon et al., 2017; de la Filia et al., 2015; Hedrick & Parker, 1997; Normark, 2003). From a theoretical perspective, most work on haplodiploidy has focused on the evolution of eusociality (Hamilton, 1964a, 1964b, 1972; Rautiala et al., 2019, but see de la Filia et al., 2015). However, haplodiploid transmission genetics can have many other important evolutionary consequences. For example, when haplodiploid populations hybridize, only female hybrids are produced in the first generation and hybrid males are produced in the subsequent generation. This asymmetry may lead to higher rates of mitochondrial introgression compared to nuclear introgression (Linnen & Farrell, 2007; Patten et al., 2015) and may have consequences for the evolution of postzygotic isolation (Bendall et al., 2020; see also Ghenu et al., 2018; Nouhaud et al., 2020). Haplodiploidy is also expected to impact the evolution of sexually selected traits (Kirkpatrick & Hall, 2004; Reeve & Pfennig, 2003), mating systems (Boulton et al., 2015; Werren, 1993), parental care (Davies & Gardner, 2014; Gardner, 2012), sex ratios (Hamilton, 1967), and the outcomes of intra- and inter-locus conflicts (Hitchcock, 2021; Kraaijeveld, 2009). However, formal theory and empirical tests for the evolutionary consequences of haplodiploidy remain rare (de la Filia et al., 2015).

Here, we focus here on how haplodiploidy affects genomic differentiation in diverging populations and species. Similarities in transmission genetics between haplodiploid genomes and X (or Z) chromosomes make it possible to draw on faster-X theory to generate predictions for haplodiploids (Avery, 1984; Hedrick & Parker, 1997; Kraaijeveld, 2009). Within populations, hemizygosity in XY and haplodiploid males will expose recessive or partially recessive mutations to selection, thereby hastening the removal of deleterious recessive alleles and the fixation of beneficial recessive alleles (Avery, 1984; Charlesworth, et al., 1987; Hedrick & Parker, 1997). More efficient selection on novel hemizygous alleles will also impact linked variation via hitchhiking (Betancourt et al., 2004) and background selection (Charlesworth, 2012), and these effects will be exacerbated by a lack of recombination in XY and haplodiploid males (Betancourt et al., 2004; Lester & Selander, 1979; Owen, 1986). So long as adaptation is driven primarily by new mutations that are at least partially recessive, faster-X theory predicts higher adaptive substitution rates and greater genetic divergence at linked sites on sex chromosomes and haplodiploid genomes relative to diploid autosomes when populations or species diverge in isolation (Presgraves 2018, but see Wright et al., 2015).

Conversely, faster-X theory under models of divergence-with-gene-flow suggests that hemizygous selection can facilitate adaptive differentiation at a selected site regardless of dominance and starting allele frequency (Lasne et al., 2017). Additionally, when formally isolated populations come into secondary contact, sex-limited hemizygosity and recombination can reduce effective migration rates at neutral loci linked to loci involved in local adaptation and/or hybrid incompatibilities (Fraïsse & Sachdeva, 2021; Fusco & Uyenoyama, 2011; Muirhead & Presgraves, 2016). So long as gene flow accompanies divergence, faster-X theory predicts that sex chromosomes and haplodiploid genomes will exhibit greater differentiation at selected and linked sites compared to autosomal chromosomes.

Consistent with faster-X theory, comparative and population genomic data from diverse taxa suggest that faster-X effects (i.e., elevated differentiation, divergence, and substitution rates on sex chromosomes) are widespread in nature (Irwin, 2018; Meisel & Connallon, 2013; Presgraves, 2018). However, these patterns are not necessarily caused by sex-limited recombination and hemizygosity. Indeed, there are many other differences between sex chromosomes and autosomes that can also produce differences in genetic differentiation, including: differences in effective population size (*N_e_*), mutation rate, recombination rate, gene content, and susceptibility to meiotic drive (Frank, 1991; Hurst & Pomiankowski, 1991; Meiklejohn et al., 2018; Patten, 2018). Because they lack sex chromosomes, haplodiploids are potentially powerful models for investigating the impact of sex-limited hemizygosity and recombination on genomic differentiation independent of sex-chromosome specific factors. However, because they also lack anything analogous to diploid autosomes, haplodiploids do not have a built-in benchmark for evaluating whether they exhibit “faster-haplodiploid” effects, which we define as greater differentiation or divergence in haplodiploids relative to comparable diploids. Fortunately, increasingly sophisticated tools for simulating genomic datasets evolving under complex demographic and ecological scenarios (Hoban et al., 2012; Haller & Messer 2019; Hart et al., 2021) offer a potential strategy for detecting faster-haplodiploid effects: simulate a benchmark diploid dataset with equivalent demographic history, recombination, and mutation under neutral and adaptive scenarios.

To better understand the impact of haplodiploidy on genomic differentiation, we combine simulations of haplodiploid and diploid genomes evolving under divergence-with-gene-flow with an empirical case study of a haplodiploid species pair for which we have extensive knowledge regarding the drivers of divergent selection and reproductive isolation, as well as basic life history knowledge to parameterize simulations. *Neodiprion pinetum* (white pine sawfly) and *N. lecontei* (redheaded pine sawfly) are sister species with overlapping distributions in eastern North American (Linnen & Farrell, 2008, 2010). Because both species are pests of economically important pines, their basic ecology and life history is well described (Benjamin, 1955; Rauf & Benjamin, 1980; Coppel & Benjamin, 1965; Knerer & Atwood, 1973; Wilson et al., 1992). Reproductive adults emerge in spring after overwintering as cocoons. Females fly to their preferred host and attract haploid males via a sex pheromone. Mating takes place on the host plant, and females use their saw-like ovipositor to embed their full complement of eggs within the needles of a single pine branch. Larvae emerge and feed on pine needles before dispersing to the soil to spin a cocoon.

While *N. pinetum* and *N. lecontei* share many similarities, *N. pinetum* feeds exclusively on white pine (*Pinus strobus*) and *N. lecontei* tends to avoid this host. Differences between their hosts likely generate divergent selection on many different larval and adult traits (Coppel & Benjamin, 1965; Codella & Raffa, 2002; Lindstedt et al., 2020; Bendall et al., 2017). For example, differences in needle chemistry and thickness between the preferred hosts of *N. lecontei* and *N. pinetum* are associated with differences in egg size, female ovipositor morphology, and female egg-laying behaviors. These traits, which together determine the reproductive success of adult females, act as an ecological barrier to gene exchange in sympatric populations (Bendall et al., 2017). This previous work suggests that many regions of the genome are likely to be under divergent selection between these species. Moreover, a coalescent-based analysis revealed evidence of historical mitochondrial introgression, suggesting that this species pair has diverged with gene flow (Linnen & Farrell, 2007).

We hypothesize that adaptation to different pines and speciation-with-gene-flow in *N. lecontei* and *N. pinetum* was facilitated by sex-limited hemizygosity and recombination. To test this hypothesis, we: (1) simulate diploid and haplodiploid “genomes” (500-kb loci) evolving under mutation, drift, divergent selection, migration, and recombination; (2) model the divergence history and characterize patterns of genomic differentiation in sympatric populations of *N. lecontei* and *N. pinetum*; and (3) use our estimated divergence history and other system-specific details to parameterize simulations of haplodiploid and diploid genomes evolving under varying levels of selection. Our data support a faster-haplodiploid effect in *Neodiprion* sawflies, and based on our results, we suggest that such effects have promoted adaptation and speciation in haplodiploid taxa.

## Methods

### Simulation of haplodiploid and diploid chromosomes under divergence with gene flow

To evaluate the effects of hemizygous selection and sex-limited recombination on genomic differentiation patterns, we simulated populations of diploid autosomes and haplodiploid chromosomes (Figure 1). We used SLiM v3 (Haller & Messer, 2019) to simulate 500-kilobase (kb) chromosomes evolving via mutation, drift, migration, and selection. We considered an isolation-with-migration model with two populations that diverged at some time (*t_div_*) from an ancestral population, with symmetric gene flow (Figure 1b). Simulations consisted of two phases. First, to enable the ancestral population to reach mutation-drift equilibrium, we simulated neutral evolution of an ancestral population with an effective size of 1500 (2*N_e_*), a mutation rate of 2.5×10^-7^/bp/generation, and a recombination rate of 2.5×10^-7^/bp/generation for 10,000 generations (>4*N_e_* generations). Second, to simulate divergence-with-gene-flow, the ancestral population splits into two equally sized populations (2*N_e_*) that exchange migrants at a constant and symmetrical migration rate (*m=m_12_=m_21_*). The timing of this split coincides with the onset of divergent natural selection on a polymorphic site (initial frequency of derived allele *a* denoted as *q_0_*) located at the middle of the chromosome (250-kb). We modeled selection under a “parallel dominance” fitness model in which the derived allele *a* is favored in population 1 and allele *A* (ancestral allele) is favored in population 2, its dominance is the same irrespective of the population (Figure 1a, as in Moran (1959); Lasne et al., (2017)). Our model assumes there are separate sexes, with equal numbers of males and females, without sex-specific migration rates, and similar distributions of offspring numbers in males and females. Our model also assumes the fitness of hemizygous males (*A* or *a*) is equal to the fitness of corresponding homozygous females (*AA* or *aa*). Following the onset of selection, populations evolve for an additional 2000 generations.

**Figure 1.**
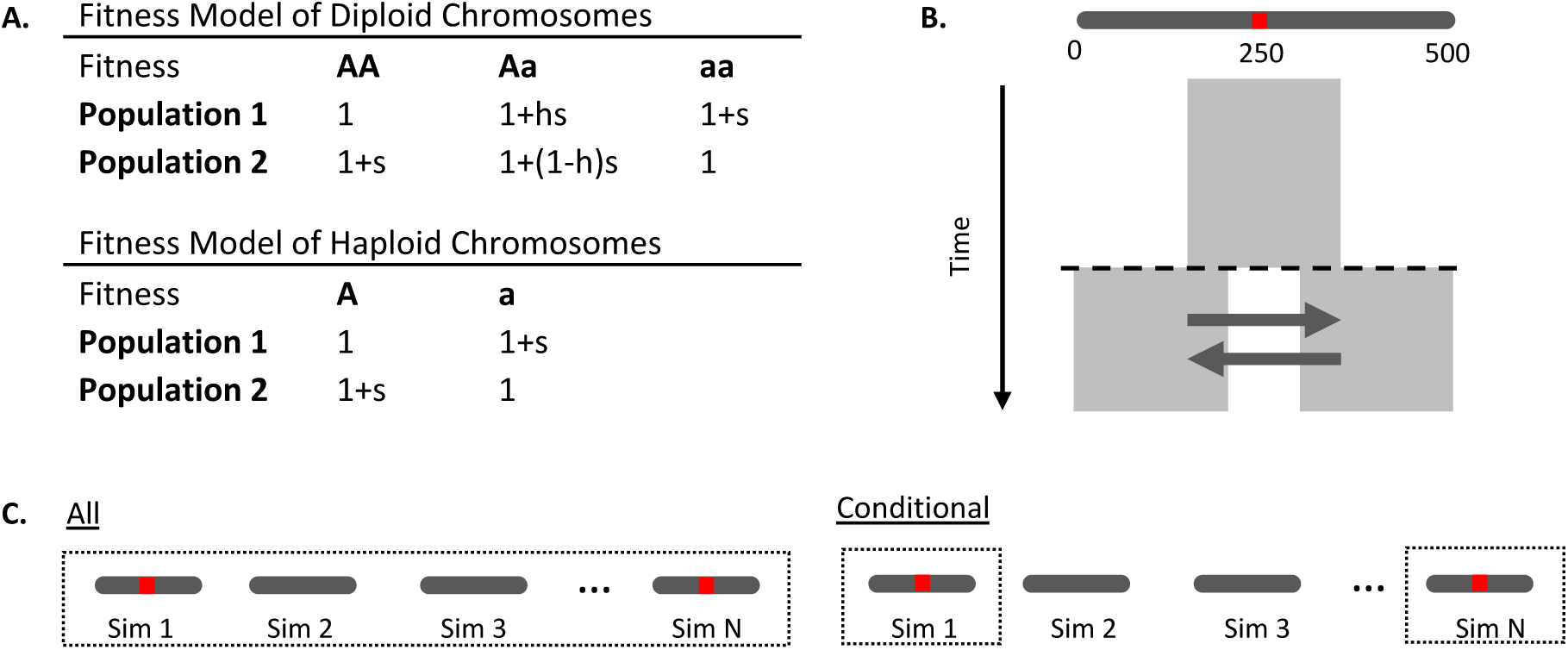
Overview of simulation approach for evaluating faster-haplodiploid effects on genomic differentiation. **A**. Fitness models for diploid chromosomes (diploid autosomes and haplodiploid females) and haploid chromosomes (haplodiploid males). This is a parallel dominance model in which the fitness of heterozygotes depends on the dominance of allele *a*, which is assumed to be the same in both populations. Population 1 is the population where the derived allele is beneficial. **B**. Overview of stochastic simulations under an isolation-with-migration model. An ancestral population with an effective size of 2*N_e_*=1500 gene copies (i.e., a haplodiploid locus with 500 females and 500 males or a diploid locus with 375 females and 375 males) of a 500-kb chromosome (dark grey bar) evolves for 10,000 generations to reach mutation-drift equilibrium. This population then splits into two equally sized populations. A divergently selected site at position 250-kb (red line) with two alleles (*A* and *a*) is introduced at the time of split, with an initial frequency *q_0_* of allele *a* in both populations. The populations evolve for *T_div_*=2000 generations, experiencing symmetric migration at a constant rate. For each parameter combination, 1000 simulations were run. **C**. Two approaches were used to summarize simulation results. In the “all”-simulations approach, mean F_ST_ is computed across all simulations (i.e., including simulations in which the derived allele *a* was lost due to drift). In the “conditional” approach, mean F_ST_ is computed only using simulations for which the derived allele *a* was retained.

The parameter values above are identical to a scaled mutation rate (*θ*=4*N_e_μL*) and recombination rate (*ρ*=4*N_e_rL*) of a 500-kb chromosome in a population with an effective size of 100,000 and a mutation rate of 2.5×10^-9^/ bp/generation. We used a smaller effective population size of 1500 and scaled the other parameters accordingly to reduce the computational burden of forward simulations, as is usually done when using SLIM (e.g., Phung et al., 2016). Our chosen divergence time correspond to a divergence with a scaled time *T_div_*/(4*N_e_*)=2/3, which is within the range of values estimated for pairs of closely related populations and species across many taxa (Hey & Pinho, 2012; Pinho & Hey, 2010), but lower than the threshold of *T_div_*/(4*N_e_*)>1 (and 2*N_e_m*<1) proposed by Hey and Pinho (2012) as a diagnostic for fully independent species.

We simulated diverging populations under all possible combinations of seven selection coefficients (scaled 2*N_e_s* ∼ 0, 10, 20, 40, 80, 100, 200), four migration rates (scaled 2*N_e_m* ∼ 0.0, 0.5, 2.5, 5.1), two dominance coefficients (recessive *h*=0.01 and codominant *h=*0.50), and four different starting allele frequencies (*q_0_* = 1/(2*N_e_*), 0.01, 0.10, and 0.50; Supplementary Table S1). These parameters were chosen to capture a range of selection coefficients and migration rates, including the neutral case (*s*=0) and the no-migration case (*m*=0). The starting allele frequencies ranged from new (*q_0_*=1/(2*N_e_*)) or rare (*q_0_*=0.01) mutations to common variants (*q_0_*=0.10 and 0.50). To investigate the impact of recombination rate on linked variation we repeated a subset of these conditions (with *q_0_*=0.10 and 0.50) under a lower recombination rate (*r*=0.1μ= 2.5×10^-8^/bp/generation). For each unique combination of parameters, we performed 1000 simulations.

To control for factors other than hemizygous selection and sex-limited recombination that might also cause differences in genomic differentiation between diploids and haplodiploids, our simulations were scaled so that haplodiploid and diploid chromosomes experience equivalent effective levels of drift, migration, and recombination (Supplementary Table S1). Thus, we adjusted the *N_e_* to ensure that both the diploid and haplodiploid chromosomes have 2*N_e_*=1500, which is the *N_e_* of a hemizygous locus with *N*=1000 individuals (500 females with two copies and 500 males with one copy). This corresponds in the diploid case to *N_D_*=750 individuals, obtained as *N_D_*=*x N* individuals, where *x* is a scaling factor that is 3/4 for 0.50 sex-ratio (Supplementary Methods). Because a haplodiploid chromosome spends 2/3 of the time in the sex in which it recombines, we scaled the diploid recombination rate as 2/3 of that of haplodiploids to ensure identical average recombination rates in diploids and haplodiploids (Kong et al., 2002; Wilfert et al., 2007). To confirm our scaling, we verified that the values of several summary statistics measuring diversity, differentiation and linkage disequilibrium were identical for neutral simulations for haplodiploid and diploid chromosomes, and that they converged to the expected values under neutrality (Supplementary Figure S1).

For each replicate, we followed the trajectories of allele frequencies at the selected site in both populations, which were used to compute the number of simulations that retained the derived allele *a* in population 1. To investigate patterns of variation in samples rather than at the population level, in the last generation we sampled 20 chromosomes of 500 kb from each population. For each parameter combination, we computed average nucleotide diversity, *D_xy_* and weighted F_ST_ across simulations using the Hudson estimator (Bhatia et al., 2013), averaged across all SNPs at three scales: (1) a 20-kb window centered on the selected site (“20-kb”); (2) across the 500-kb chromosome (“500-kb”); and (3) scan of contiguous non-overlapping 20-kb windows (“genome scan”). To measure patterns of linkage disequilibrium, we computed average *r^2^* between all pairs of SNPs within 20-kb windows. To evaluate the effect of loss of the derived allele *a*, we used two approaches: in the “all simulations” approach, the mean of summary statistics (e.g., F_ST_) was computed across all simulations, whereas in the “conditional” approach, the mean of summary statistics is computed only across simulations where the derived allele was retained in population 1 (Figure 1c). Thus, the “conditional” approach removes the effect of allele loss. Parameter combinations for which fewer than 10 simulations were retained were treated as missing data.

### Empirical data: estimating divergence history in a haplodiploid species pair

#### Population sampling

We sampled 23 *N. pinetum* larvae and 44 *N. lecontei* larvae from Kentucky (Supplementary Table S2). Larvae tend to be found in gregarious colonies of siblings in both species. To ensure we were not sampling close relatives, each individual was collected from a different colony. To maximize our chances of sequencing diploid female larvae, which tend to be larger than haploid male larvae, we extracted DNA from large larvae and verified sex with heterozygosity estimates. We also sampled 3 suspected field-caught hybrids (based on intermediate larval pigmentation) from Kentucky and one lab-reared female F_1_ as a control (Supplementary Table S2). An additional 18 *N. lecontei* samples from an allopatric population in Michigan and 1 *N. virginiana* from Blackstone, VA (37°06’47.2”N, 78°01’37.4”W) were sequenced for use in demographic analyses.

#### DNA sequencing

We extracted DNA using a CTAB/phenol-chloroform-isoamyl alcohol method (Chen, Rangasamy, Tan, Wang, & Siegfried, 2010). We visualized the DNA on a 0.8% agarose gel to confirm quality. To quantify the DNA, we used a Quant-iT High-Sensitivity DNA Assay Kit (Invitrogen – Molecular Probes, Eugene, OR, USA). For *N. pinetum*, *N. lecontei*, and hybrids, we used a modified ddRAD sequencing protocol from Bagley et al. (2017) and Peterson et al. (2012). We fragmented the DNA using NlaIII and EcoRI. We assigned each individual along with additional samples from other projects to one of eight libraries. During adapter ligation, we assigned each sample one 48 unique in-line barcodes (Supplementary Table S2). We used the 5-10 bp variable length barcodes used in Burford Reiskind et al. (2016). We then pooled each group of samples and size selected for a 379-bp fragment (+/- 76bp) on a PippinPrep (Sage Science, Beverly, MA). We did 12 rounds of high-fidelity PCR amplification (Phusion High-Fidelity DNA Polymerase, NEB, Ipswich, MA) using PCR primers that included one of 12 unique Illumina multiplex read indices (Supplementary Table S2). To allow for the detection of PCR duplicates, we included a string of 4 degenerate bases next to the Illumina read index (Schweyen et al., 2014). We used a Bioanalyzer 2100 (Agilent, Santa Clara, CA) to check library quality. The libraries were sequenced at the University of Illinois’ Roy J. Carver Biotechnology Center, using two lanes of Illumina HiSeq 4000 and150-bp single-end reads.

For *N. virginiana*, which we used as an outgroup, we used 150-PE reads generated on an Illumina Nextseq at the Univeristy of Georgia Genomics Facility (Vertacnik, 2020). Library preparation and whole-genome shotgun sequencing were both completed at the sequencing facility. We removed the adapters using cutadapt 1.16 and contaminants using the standard and pine databases in Kraken (Martin, 2011; Wood & Salzberg, 2014).

#### DNA processing and variant calling

We aligned demultiplexed ddRAD reads to the *N. lecontei* reference genome (Nlec1.1 GenBank assembly accession number- GCA_001263575.2; Linnen et al., 2018; Vertacnik & Linnen, 2015) using the very sensitive setting in bowtie2 (Langmead & Salzberg, 2012). We only retained reads that aligned to one locus in the reference genome and had a Phred score greater than 30. For the ddRAD dataset, we removed PCR duplicates using a custom script. We called SNPs in samtools (Li et al., 2009). Male and female larvae are morphologically indistinguishable. To identify putative haploid males, which are expected to have unusually low heterozygosity, we computed per-individual heterozygosity (as in Bagley et al., 2017). No individuals were excluded based on heterozygosity. We required all sites to have a minimum of 7x coverage and 50% missing data or less. We also removed SNPs with significantly more heterozygotes than expected under Hardy-Weinberg equilibrium (an indicator of genotyping/mapping error). We removed any individual that was missing more than 70% of the data. We performed all filtering in VCFtools v0.1.13 (Danecek et al., 2011).

We created several datasets with subsets of individuals and additional filtering for each of the population genetic analyses. We generated three data sets with minor allele filtering (MAF, SNPs <0.01 removed): 1) sympatric *N. pinetum* and *N. lecontei* for genome-wide patterns of divergence (36,935 SNPs), 2) sympatric *N. pinetum*, *N. lecontei,* and hybrids for admixture analysis (35,649 SNPs), and 3) sympatric *N. pinetum*, *N. lecontei,* allopatric *N. lecontei*, and outgroup *N. virginiana* for ABBA-BABA tests (12,905 SNPs). We also generated a down-sampled dataset (described below) without a MAF filter for estimating site-frequency spectra (SFS) that included sympatric *N. pinetum*, *N. lecontei*, and *N. virginiana* for demographic analyses.

#### Population structure, demographic analysis, and genomic differentiation

To confirm that our morphological hybrids were genetically admixed, we used Admixture v1.3.0 (Alexander et al., 2009) to estimate the proportion of ancestry for each individual collected in Kentucky (*N. lecontei, N. pinetum,* lab-reared hybrids, and suspected field-caught hybrids) from *K* populations for *K* =1-5. We ran 100 replicates per *K* and chose the *K* with the lowest cross-validation (CV) score (Supplementary Table S3). Additionally, to test for introgression between sympatric *N. lecontei* (P1) and *N. pinetum* (P3), we performed an ABBA-BABA test (Patterson et al., 2012) with Kentucky *N. lecontei* (P1), Michigan *N. lecontei* (allopatric population, P2), Kentucky *N. pinetum* (P3) and *N. virginiana* (outgroup, P4). We used custom R script to compute the ABBA-BABA assuming that the outgroup is not fixed for the ancestral allele (Patterson et al., 2012), assessing significance with block-jackknife resampling diving data into 645 blocks of ∼20 SNPs.

To evaluate the timing and magnitude of gene flow between *N. lecontei* and *N. pinetum*, we performed demographic modeling based on the site frequency spectrum (SFS) using the composite likelihood method implemented in fastsimcoal2 v2.6 (Excoffier et al., 2013). For this analysis, we used ddRAD data from sympatric populations of *N. lecontei* and *N. pinetum* filtered as described above, with additional filters applied to satisfy analysis assumptions. First, to minimize the impact of selection on demographic history estimates, we used the NCBI *Neodiprion lecontei* Annotation Release 100 (updated to GCA_001263575.2) to exclude SNPs that were in or within 1 kb of the start or end of a gene, thereby generating a set of putatively neutral markers. Furthermore, to reduce bias in the SFS, we applied more stringent depth-of-coverage filters, requiring a minimum depth of 10x and a maximum depth less than 2x the median depth of coverage per individual. To build the 2D-SFS without missing data, each scaffold was divided into non-overlapping 50-kb blocks, and we kept only blocks where the median distance between SNPs was larger than 2bp. SNPs without missing data were obtained for each block by downsampling four and six females from *N. pinetum* and *N. lecontei*, respectively. This resulted in a downsampled dataset with 9,994 SNPs. To polarize the ancestral/derived state of alleles and obtain the unfolded 2D-SFS we used data from *N. virginiana*. To obtain the number of invariant sites in the 2D-SFS we assumed that the proportion of SNPs removed because of extra filters was the same for invariant sites. Given a proportion of number of SNPs to number of invariant sites before extra filters of ∼0.046, the number of invariant sites in the 2D-SFS after filters was set to 215,283.

We tested five alternative demographic scenarios: (1) divergence without gene flow, (2) divergence with continuous bidirectional migration, (3) divergence in isolation followed by a single bout of secondary contact (unidirectional gene flow), (4) divergence with bidirectional migration that stops before divergence is complete, and (5) divergence in isolation followed by continuous secondary contact (bidirectional). All models except the model of continuous gene flow had an equal number of parameters, so we compared their likelihoods directly. We ran each model 100 times starting from different parameter combinations, each run with 50 optimization cycles (-l50) and approximating the expected SFS with 100,000 coalescent simulations (- n100000). We selected the run with the highest likelihood to estimate parameter values.

To examine genome-wide patterns of genetic divergence, we computed F_ST_ and π in 100- kb non-overlapping windows for *N. lecontei* and *N. pinetum* in VCFtools on the non-down sampled dataset. To identify regions of the genome that were more or less differentiated than expected under neutrality, we simulated 10,000 datasets under the inferred demographic history for sawflies using coalescent simulations implemented in the R package *scrm* (Staab et al., 2015). For each simulation, we computed F_ST_ as for the *Neodiprion* dataset (see above). Outlier windows were defined as those above or below the 95% CI for F_ST_ obtained from the 10,000 simulations. Simulations were done assuming no recombination, 50% missing data (i.e., female samples sizes of 0.5*23 for *N. pinetum* and 0.5*44 for *N. lecontei*), and scaling theta (*4Nμ*) such that the average number of SNPs per window across simulations were similar to the observed in *Neodiprion* dataset.

### Comparison of empirical haplodiploid data to simulated diploids and haplodiploids

To evaluate the potential influence of haplodiploidy on genomic differentiation between *N. lecontei* and *N. pinetum*, we used SLiM v3 to simulate haplodiploid and diploid “genomes” evolving under divergent selection and the demographic history estimated for our focal species pair. Because *N. pinetum* is on the derived host plant (Linnen & Farrell, 2010), we modeled *N. pinetum* as population 1 (where derived allele *a* is favored), and *N. lecontei* as population 2. For these simulations, we also assumed a sex ratio of 70 females:30 males based on previously published sex ratios for *N. lecontei* and *N. pinetum* (Harper et al., 2016). As in our first set of simulations, we scaled our simulations to ensure equivalent levels of drift, migration, mutation rates and recombination between diploids and haplodiploids.

To reduce the computational burden of forward SLiM simulations we re-scaled parameters such that under neutrality, the SFS obtained with SLiM was identical to the expected SFS obtained under the demographic history inferred with fastsimcoal2. This was achieved by ensuring that the scaled mutation rate 4*N_A_μL* for *L* sites was identical in both cases, where *N_A_* is the ancestral effective size and *μ* is the mutation rate per site per generation. By considering a mutation rate two orders of magnitude higher (*μ*=3.50×10^-7^ rather than the 3.50×10^-9^/bp/generation used for SFS-based inference) and that *L*=5×10^5^ sites in SLiM corresponds to *L*=5×10^4^ sites in the 2D-SFS used for fastsimcoal2 (including SNPs and invariant sites), the haploid effective population sizes were three orders of magnitude lower (328 for *N. pinetum*, 1093 for *N. lecontei* and 1982 for the ancestral population) and migration rates three orders of magnitude higher (3.64×10^-4^ into *N. pinetum*, 1.71×10^-5^ into *N. lecontei*). The times of split were scaled accordingly, resulting in 1548 generations (rather than 1.54×10^6^). To obtain the number of individuals *N* in SLiM that correspond to the above haploid effective sizes, we had to account for the sex-ratio of 70 females:30 males (Supplementary Methods). Given the average of 19.02 SNPs in *Neodiprion* 100-kb windows (with gaps due to sparse ddRAD loci), the average number of SNPs in SLiM simulations under neutrality of 2,257 would correspond to ∼10Mb of a similar ddRAD dataset. We considered two recombination rates *r*: (i) equal to the mutation rate (*r*=*μ*=3.50×10^-7^), and (ii) three times higher than mutation rate (*r*=1.05×10^-6^) using an estimate of 3.43 cM/Mb based on a linkage map for *N. lecontei* constructed from an inter-population cross (Linnen et al., 2018). In both cases, recombination rate for diploids was scaled by (2/3) as done for the simulation study (see above). Because the per-locus selection estimate is unknown, we simulated differentiation and under a wide range of selection coefficients *s* from 0.0 to 0.3. We also simulated all combinations of two dominance coefficients (*h* = 0.01 and 0.50) and one starting allele frequency (*q_0_* = 0.10) (Supplementary Table S4). We computed mean F_ST_ across all 1000 simulations for each starting allele frequency, dominance, and selection coefficient combination. These combined simulations can be thought of as a divergence history in which, on average, there is a divergently selected site every 10Mb.

## Results

### Faster-haplodiploid effects on genomic differentiation with migration

Across all parameter combinations and for both window sizes (20-kb and 500-kb), we found that F_ST_ between haplodiploid populations was always equal to or greater than F_ST_ between diploid populations (Figure 2). Migration was required for haplodiploid F_ST_ to exceed diploid F_ST_. For both window sizes, we found that faster-haplodiploid effects (i.e., ratio of haplodiploid F_ST_: diploid F_ST_ > 1) were more pronounced in the recessive case (Figure 2a-d) than in the codominant case (Figure 2e-h). For each dominance coefficient, the regions of parameter space that *maximized* faster-haplodiploid effects depended on the window size used to calculate F_ST_. For sites tightly linked to the selected site (20-kb window), faster-haplodiploid effects were maximized when migration was high (2*N_e_m≥*2.5) and selection was moderate (10≤2*N_e_s≤*40) (Figures 2b and 2f). By contrast, when we considered much larger windows of 500-kb, relative differences between haplodiploid and diploid differentiation levels were maximized at higher selection coefficients (2*N_e_s≥*40) but at similar migration rates (2*N_e_m≥*2.5). The same trend—faster-haplodiploid effects maximized at higher selective coefficients for the 500-kb windows than for the 20-kb windows—was found for all initial frequencies of the derived allele *a* for the codominant case and for *q_0_=*0.5 for the recessive case (Supplementary Figure S2). However, for new (*q_0_=*1/(2*N*)) or rare (*q_0_=*0.01) recessive alleles, faster-haplodiploid effects were always maximized at the highest selection coefficients, regardless of window size (Supplementary Figure S2).

**Figure 2.**
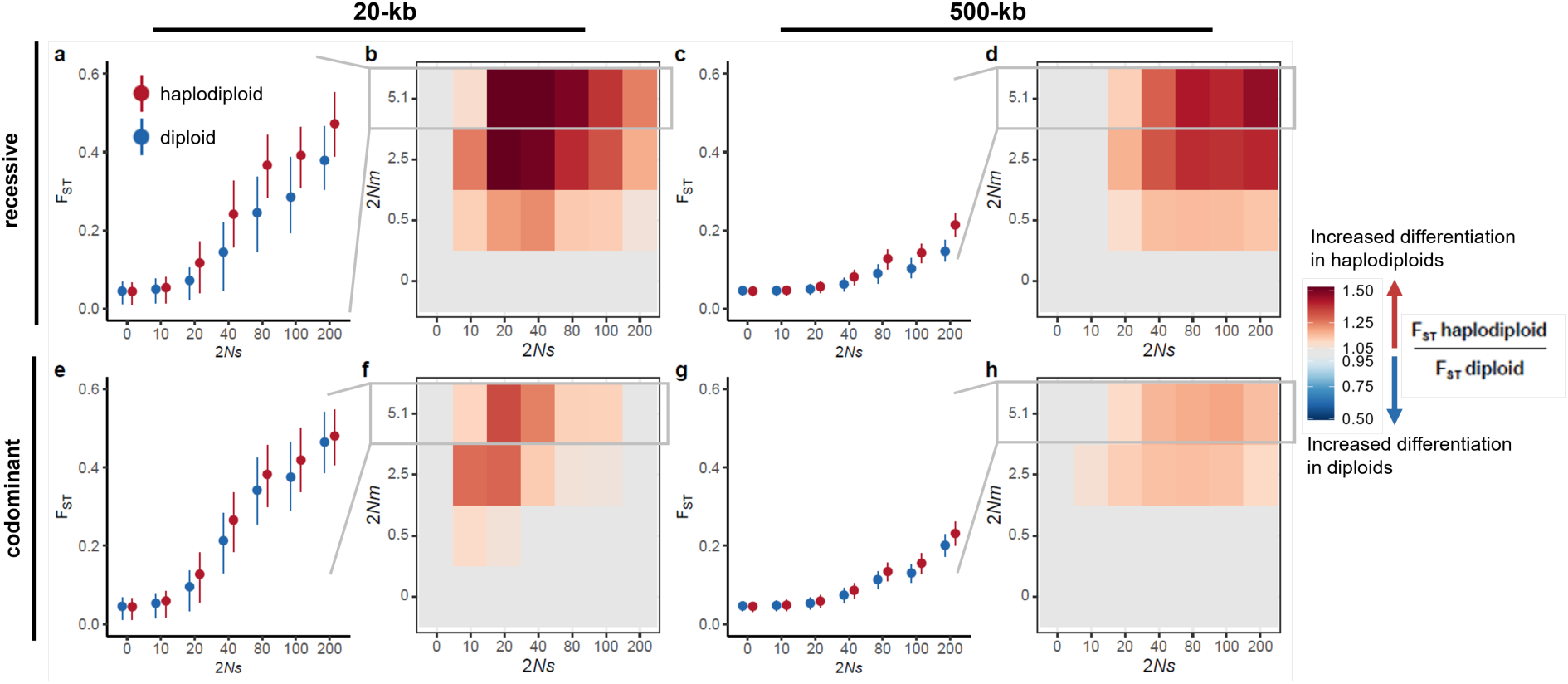
Faster-haplodiploid effects as a function of strength of divergent selection, migration rate and dominance. **(a, e, c, g)** Differentiation (F_ST_) for haplodiploids and diploids with *2Nm*=5.1 and varying selective coefficients (*2Ns*) for different window sizes with selected site in the middle [**(a, e)** 20-kb, **(c, g)** 500-kb], and different dominance coefficients [**(a, c)** recessive (*h*=0.01) and **(e, g)** codominant (*h*=0.50)]. The points correspond to mean F_ST_ and the whiskers to the interquantile range based on 1000 simulations. **(b, d, f, h)** Heatmap of the ratio of haplodiploid to diploid (H/D) mean F_ST_ for different combinations of selective coefficients and migration rates for different window sizes [**(b, f)** 20-kb and **(d, h)** 500-kb], and dominance coefficients [**(b, d)** recessive (*h*=0.01) and **(f, h)** codominant (*h*=0.50)]. Results were obtained with 1000 simulations for each parameter combination with an initial frequency *q_0_*=0.10, sampling 20 females from each population.

One mechanism leading to greater differentiation in haplodiploids was differential allele loss. Simulations of haplodiploid populations had a higher probability of retaining the derived allele across a wide range of parameter combinations (Figure 3, Supplementary Figure S3). For recessive alleles, differences in allele retention between haplodiploids and diploids were dependent on both starting allele frequency and selection strength, but relatively insensitive to migration rate (Figure 3a; Supplementary Figure S3). By contrast, outside of a slightly elevated probability of retaining the derived allele at lower selection coefficients (10 *≤* 2*N_e_s ≤* 20, inset panel in Figure 3b), differences in allele retention between haplodiploid and diploid populations were minimal for the codominant case (Figure 3b). For both recessive and codominant cases, increasing migration had a limited but consistent effect, leading to a lower probability of allele retention.

**Figure 3.**
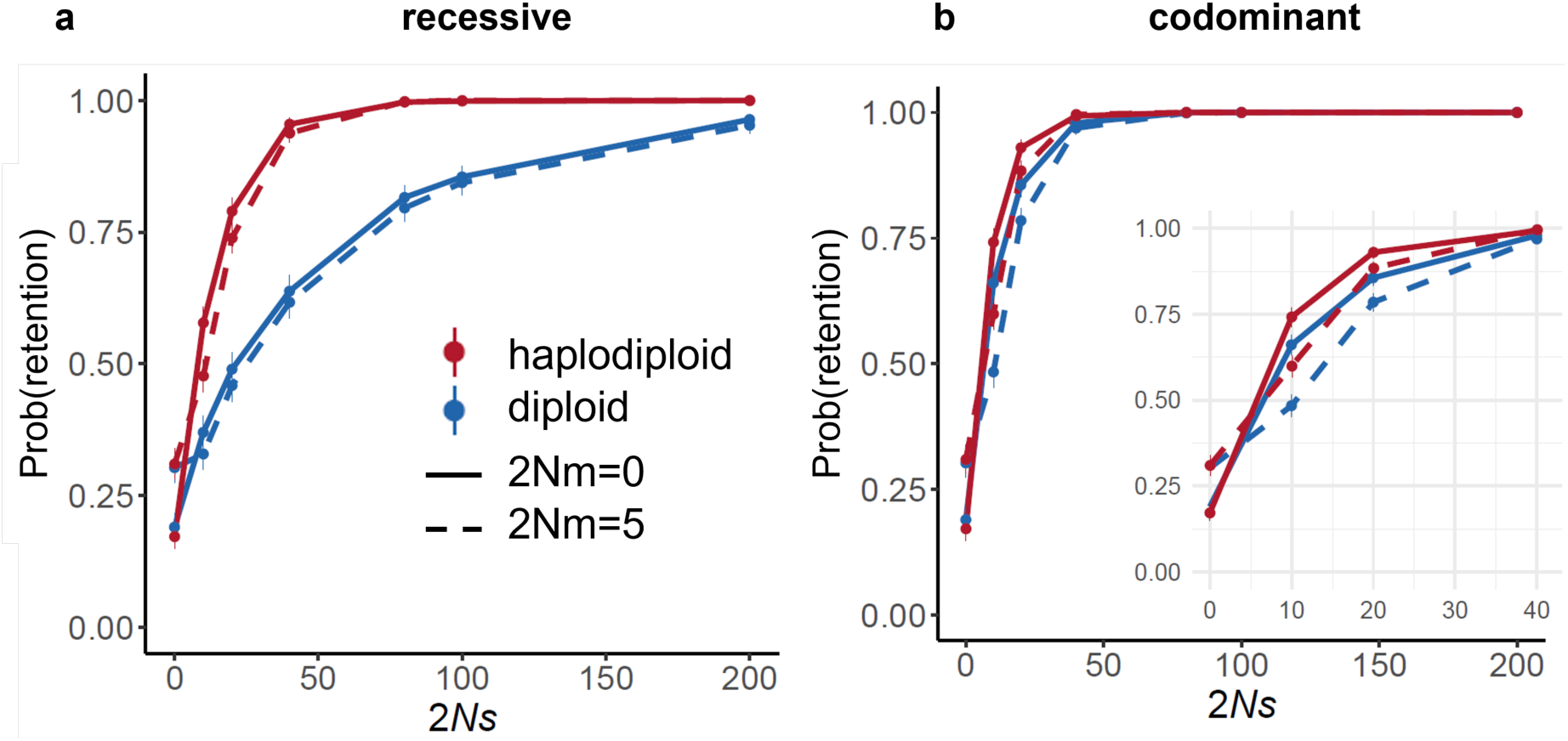
Effect of haplodiploidy on retaining the derived allele at the site under divergent selection. Probability of retaining - Prob(retention) - the derived allele *a* for haplodiploid and diploid simulations at the two extremes of migration rates considered: no migration (2*Nm*=0; solid lines) and high migration (2*Nm*=5.1, dotted lines), for **a)** recessive (h=0.01), and **b)** codominant (h=0.50) mutations. In **b)**, inset shows a zoom for *2Ns* values between 0 and 40. Probability of allele retention is calculated as the proportion out of 1000 simulations that retained the derived allele *a* in population 1 (where *a* is favored). 95% confidence intervals are Clopper-Pearson CI for proportions. These results are for an initial allele frequency of 0.10 (*q_0_*=0.10).

To investigate the impact of haplodiploidy on genomic differentiation via mechanisms other than differential allele loss, we computed average F_ST_ for each parameter combination conditional on retaining the derived allele at the selected locus. For the recessive case, controlling for the impact of differential allele loss decreased the magnitude of faster-haplodiploid effects across all parameter combinations (Fig. 4a vs Fig. 2b and Fig. 4e vs. Fig. 2d), indicating that the increased retention of the derived allele in haplodiploids contributed to faster-haplodiploid evolution. Compared to the recessive case, conditioning on allele retention had much less of an impact on the magnitude of faster-haplodiploid effects for the codominant case. The heatmaps in Figure 5a and 5e are nearly identical to those in Figure 2f and 2h, respectively. This is unsurprising since differences in allele loss/retention were minimal in the codominant case (Figure 3). Finally, once we conditioned on retaining the derived allele, starting allele frequency had little impact on patterns of faster-haplodiploid evolution (Supplementary Figure S4; for comparison see Supplementary Figure S2).

**Figure 4.**
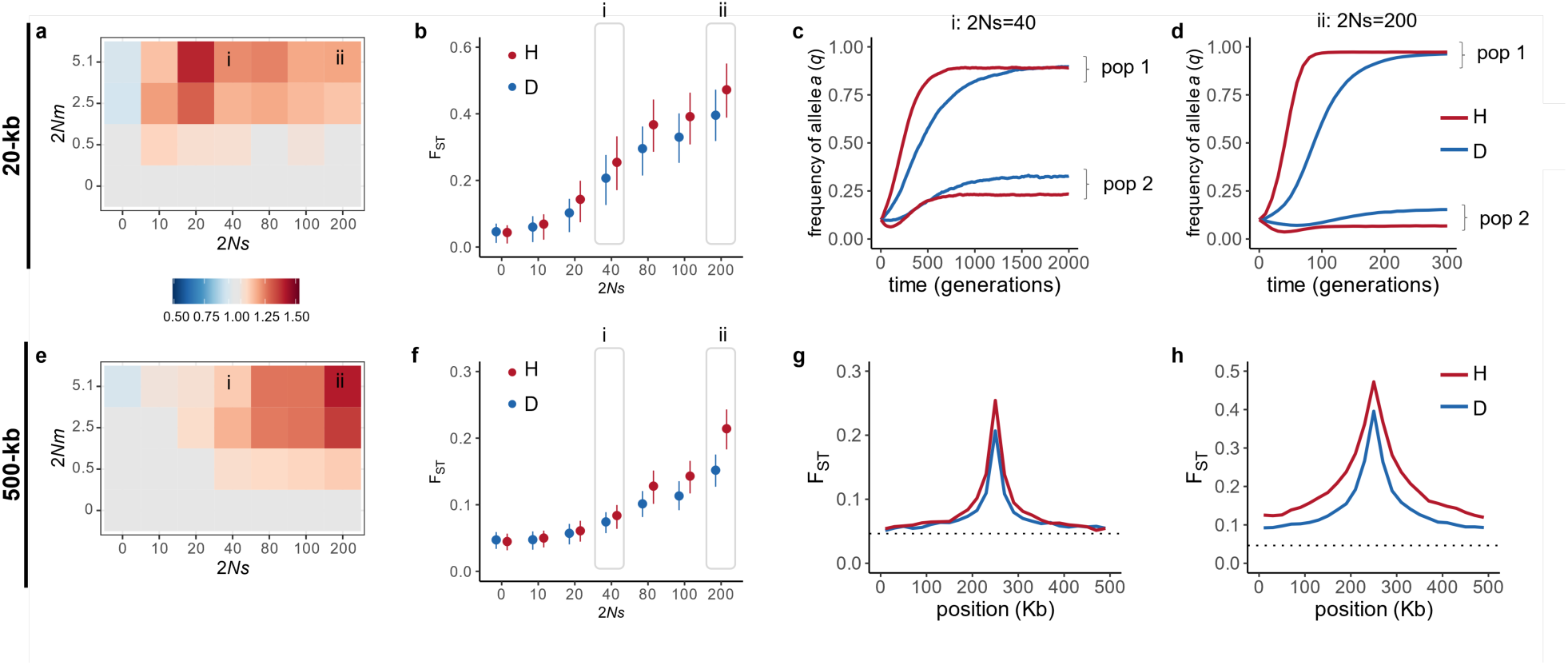
Faster-haplodiploid effects for a recessive (*h*=0.01) derived allele after removing the effect of differential allele loss. **(a, e)** Heatmap of ratio of haplodiploid (H) to diploid (D) mean F_ST_ for a combination of selective coefficients and migration rates for different window sizes with the selected site at the middle: **(a)** 20-kb and **(e)** 500-kb. Labels **i** and **ii** indicate the two cases selected to illustrate allele trajectories and scans of differentiation. **(b, f)** F_ST_ for haplodiploids and diploids with high migration (2*Nm*=5.1) and varying selective coefficients (2*Ns*), for different window sizes: **(b)** 20-kb and **(f)** 500-kb. Points correspond to mean F_ST_ and whiskers to interquantile ranges. Labels **i** and **ii** indicate the two cases selected to illustrate allele trajectories and scans of differentiation. **(c, d)** Trajectories of allele frequencies at the site under divergent selection in both populations, for: **(c)** moderate selection (2*Ns*=40), and **(d)** strong selection (2*Ns*=200). Note that the time scale is different because equilibrium differentiation is reached faster under strong selection. **(g, h)** Scan of mean F_ST_ along the 500-kb chromosome in non-overlapping 20-kb windows, obtained for: **(g)** moderate selection (2*Ns*=40), and **(h)** strong selection (2*Ns*=200). Mean and interquantile F_ST_ are based on the simulations out of 1000 that kept the derived allele *a* in population 1 at the site under divergent selection. Results are for simulations with an initial frequency *q_0_*=0.10, sampling 20 females from each population.

**Figure 5.**
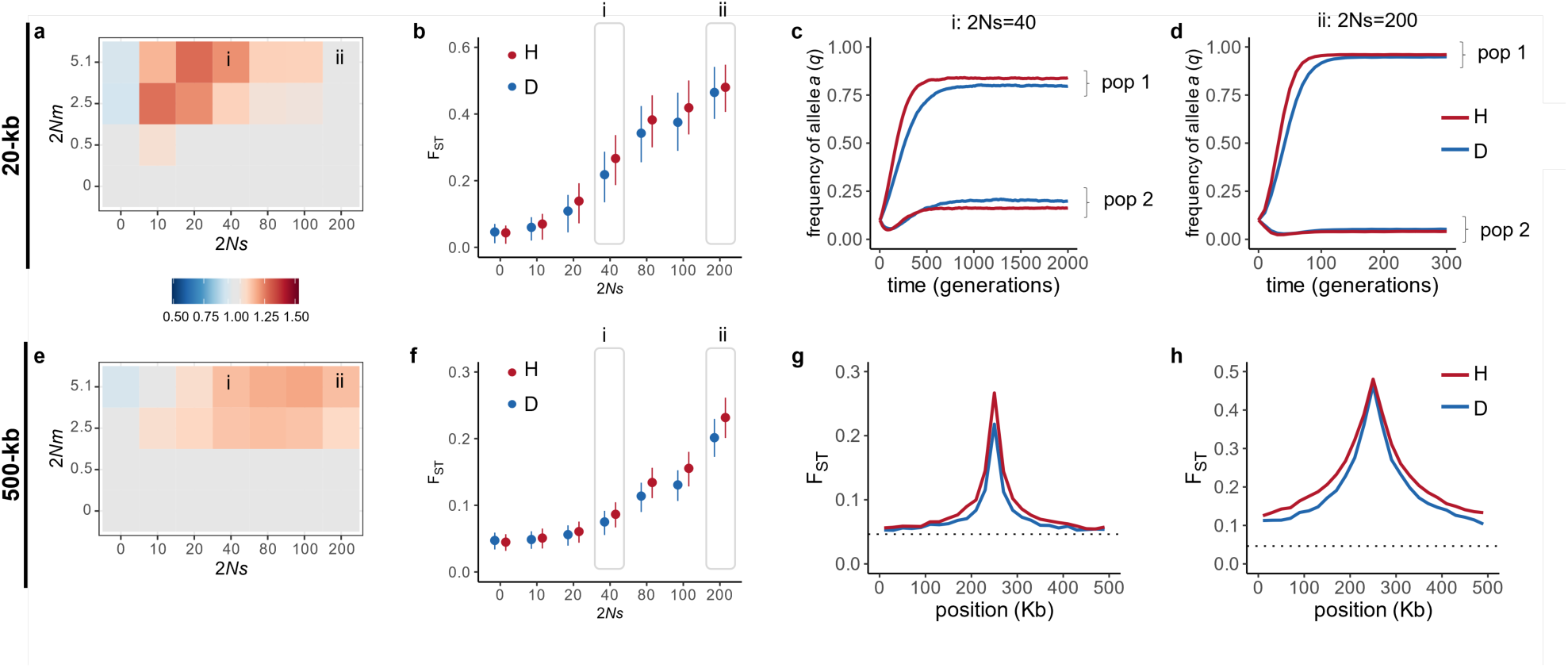
Faster-haplodiploid effects for a codominant (*h*=0.50) derived allele after removing the effect of differential allele loss. **(a, e)** Heatmap of ratio of haplodiploid (H) to diploid (D) mean F_ST_ for a combination of selective coefficients and migration rates for different window sizes with the selected site at the middle: **(a)** 20-kb and **(e)** 500-kb. Labels **i** and **ii** indicate the two cases selected to illustrate allele trajectories and scans of differentiation. **(b, f)** F_ST_ for haplodiploids and diploids with high migration (2*Nm*=5.1) and varying selective coefficients (2*Ns*), for different window sizes: **(b)** 20-kb and **(f)** 500-kb. Points correspond to mean F_ST_ and whiskers to interquantile ranges. Labels **i** and **ii** indicate the two cases selected to illustrate allele trajectories and scans of differentiation. **(c, d)** Trajectories of allele frequencies at the site under divergent selection in both populations, for: **(c)** moderate selection (2*Ns*=40), and **(d)** strong selection (2*Ns*=200). Note that the time scale is different because equilibrium differentiation is reached faster under strong selection. **(g, h)** Scan of mean F_ST_ along the 500-kb chromosome in non-overlapping 20-kb windows, obtained for: **(g)** moderate selection (2*Ns*=40), and **(h)** strong selection (2*Ns*=200). Mean and interquantile F_ST_ are based on the simulations out of 1000 that kept the derived allele *a* in population 1 at the site under divergent selection. Results are for simulations with an initial frequency *q_0_*=0.10, sampling 20 females from each population.

The observation that haplodiploid F_ST_ tends to exceed diploid F_ST_ even after conditioning on retaining the derived allele (Fig. 4a,b,e,f; Fig. 5a,b,e,f) indicates that mechanisms other than differential allele retention must contribute to elevated differentiation in haplodiploids. To gain intuition about these mechanisms note that with migration, there is a constant influx of maladapted alleles into each population. For diploids, hybridization between immigrants and residents is likely to generate hybrids that are heterozygous for the locally adaptive allele (*Aa*). The heterozygotes (*Aa*) are important because the fate of maladapted alleles is determined by differences in fitness between heterozygotes (*w_Aa_*) and resident homozygotes (*w_aa_* in population 1, *w_AA_* in population 2)(Yeaman & Otto, 2011). Additionally, the fate of linked neutral sites ultimately depends on recombination in *Aa* heterozygotes. For haplodiploids, there are no heterozygote males (Figure 1a), and males do not recombine, which likely affects (i) how efficiently selection removes maladaptive alleles at the selected site; and (ii) how recombination breaks immigrant haplotypes allowing for gene flow at neutral sites that escape linkage to maladapted alleles. The impact of the lack of heterozygous males in haplodiploids can be determined by comparing haplodiploid and diploid allele frequency trajectories at the selected site (which reveals differences in the efficacy of selection against maladapted alleles) and F_ST_ scans along the 500-kb chromosome (which reveals differences in effective migration rate at linked sites).

We examined allele trajectories and chromosome-wide F_ST_ patterns under high migration (2*Nm*=5.1) and two selection intensities (moderate: 2*Ns*=40 and strong: 2Ns=200). For sites tightly linked to recessive alleles (20-kb windows), elevated haplodiploid differentiation was observed for both selection coefficients (Figure 4b). Simulated allele trajectories at the site under divergent selection reveal two possible sources of elevated differentiation. First, the derived allele increased in frequency in population 1 in the haplodiploids more rapidly than in the diploids (Figure 4c,d). Second, once an equilibrium between migration and selection was reached, selection was more efficient at removing maladapted immigrant recessive *a* alleles in population 2 in the haplodiploids. In population 2, haplodiploid *A* males had a higher fitness than immigrant *a* males (*w_A_ > w_a_*); whereas the fitness of diploid *AA* males was similar to *Aa* males (*w_AA_ ≈ w_Aa_*, Figure 1a). As a result, the frequency of *a-*bearing haplotypes was lower in population 2—and F_ST_ between populations was higher—in haplodiploids. We attribute these differences in allele trajectories to differences in male fitness, since haplodiploid and diploid females have the same fitness (Figure 1a) and we assumed a 50:50 sex-ratio with no sex-specific migration rates.

For codominant and tightly linked sites (20-kb window), a faster-haplodiploid effect was more pronounced for moderate selection than for strong selection (Figure 5a,b). This can be explained using as an approximation for the efficiency of selection, the fitness difference between heterozygotes and resident males for diploids (*Δw_d_*=*w_AA_ – w_Aa_*=*s*/2), and between immigrant and resident males for haplodiploids (*Δw_h_* =*w_A_ – w_a_* =*s*, Figure 1a). Our results are consistent with this approximation, showing that even for the codominant case, selection is more efficient in haplodiploids (*s > s*/2). Under strong selection, by contrast, locally adaptive alleles were nearly fixed in both populations (Figure 5d). Because there were few heterozygotes in either population, differences in the efficacy of selection in haplodiploid and diploid males had a negligible impact on differentiation at the selected site.

For 500-kb windows, a faster-haplodiploid effect was more pronounced for strong selection than for moderate selection for both recessive (Figure 4e-f) and codominant (Figure 5e-f) cases. At moderate selection coefficients, differentiation across the 500-kb region was elevated at the selected site and nearby sites but dropped off quickly to the expected neutral F_ST_ (Figure 4g, 5g). This is consistent with moderately strong divergent selection reducing the effective migration rate only at neutral sites tightly linked to the selected site. By contrast, under strong selection, chromosome-wide differentiation was elevated above neutral expectations for both haplodiploids and diploids, even at distant sites (Figure 4h, 5h). We attribute the elevated differentiation across the whole 500-kb region in haplodiploids to reduced opportunities for recombination between divergently selected haplotypes. Such recombination events ultimately depend on the frequency of *Aa* heterozygotes, and for haplodiploids, a heterozygote *Aa* can only occur and recombine in females.

Our results reveal that haplodiploidy reduced opportunities for recombination in heterozygotes at two distinct phases. First, during the initial stages of divergence, haplodiploids reached migration-selection equilibrium faster than diploids: approximately ∼3x faster for the recessive case (Figure 4c-d) and ∼1.2x faster for the codominant case (Figure 5c-d). The faster time to equilibrium in the codominant case resulted in a chromosome-wide faster-haplodiploid effect even though there was no difference in F_ST_ at the selected site (Figure 5a,e). Second, once migration-selection equilibrium was reached, the frequency of the maladapted allele in each population tended to be lower in haplodiploids than diploids (Figures 4c-d, 5c-d), resulting in a lower frequency of *Aa* heterozygotes and a smaller effective recombination rate. This was especially pronounced in the recessive case. As expected, simulations with a lower recombination rate caused faster-haplodiploid differentiation of 500-kb to more closely resemble the 20-kb window results (Supplemental Figure S5).

Overall, our simulations suggest so long as there is migration, haplodiploidy will lead to elevated differentiation at selected sites and linked neutral sites when populations diverge via divergent selection. This “faster-haplodiploid effect” is produced under a wide range of selection coefficients; regardless of whether selection acts on new mutations, rare standing genetic variation, or common standing genetic variation; and regardless of whether selection acts on recessive or codominant alleles (Figure 2, Supplementary Figure S2). Finally, while the effects of hemizygous selection and sex-limited recombination tend to be most pronounced at selected sites and tightly linked neutral sites (20-kb windows), faster-haplodiploid effects can extend far beyond the selected site (500-kb windows, which translates to ∼4.4 cM in our simulations).

### Demography and genomic differentiation in pine sawflies

*Neodiprion lecontei* and *N. pinetum* differ in many host-related traits (Figure 6A). Our admixture analysis of *N. lecontei*, *N. pinetum*, a lab-reared F_1_ hybrid, and three suspected wild-caught hybrids supported two distinct genetic clusters (K=2; Supplementary Table S3). Putative wild hybrids were indistinguishable from the lab-reared hybrid, and all four individuals were genetically admixed with approximately equal contributions from *N. pinetum* and *N. lecontei* (Figure 6B). The only other admixed individual detected was morphologically indistinguishable from *N. pinetum*, but an estimated ∼13% of its genome came from *N. lecontei*. In addition to finding evidence of recent admixture, an ABBA-BABA test revealed evidence of historical introgression between sympatric *N. pinetum* and *N. lecontei* populations (*D*=0.18; *P* = 2.12 x 10^- 15^). Finally, demographic models that had no migration or only a single burst of admixture were far less likely than models that included continuous migration (Table 1). Together, these results support a divergence-with-gene-flow scenario for *N. lecontei* and *N. pinetum* (Figure 6C).

**Figure 6.**
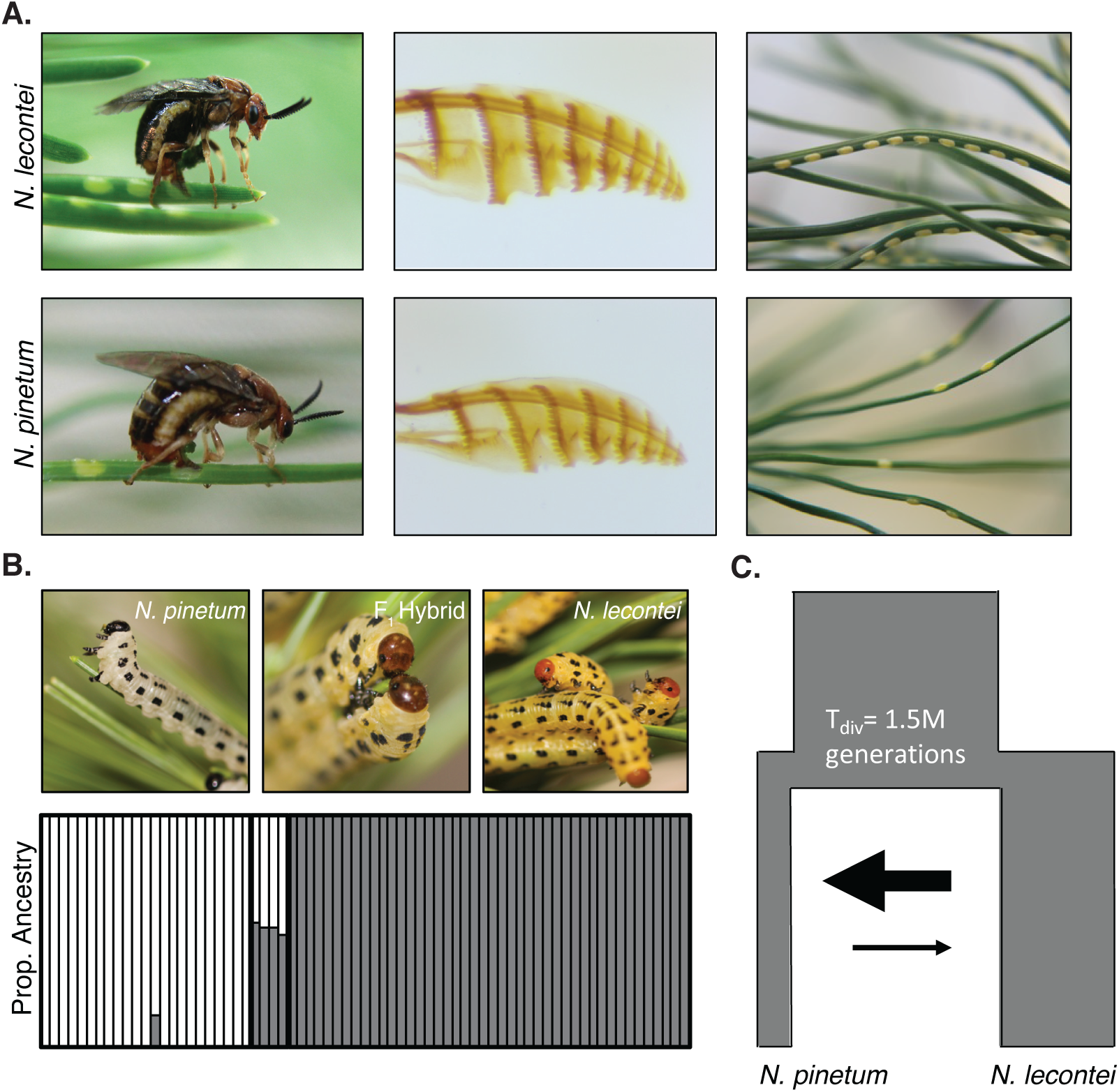
Divergent selection and divergence-with-gene-flow in pine sawflies (*N. pinetum* and *N. lecontei*). **A.** *N. pinetum* and *N. lecontei* differ in multiple oviposition traits, such as host preference, oviposition stance, ovipositor morphology, and oviposition pattern resulting in strong extrinsic postzygotic isolation. These oviposition traits along with additional host-use adaptations likely result in multiple independent regions of the genome experiencing divergent selection. **B.** Representative pictures of *N. pinetum*, *N. lecontei*, and F_1_ hybrid larvae above an Admixture plot (K=2) of individuals sampled from Kentucky. *N. pinetum* ancestry is in white; *N. lecontei* ancestry is in grey. Lab-reared (N=1) and field-caught (N=3) hybrids are genetically admixed with approximately half of their ancestry coming from each species. **C.** *N. pinetum* and *N. lecontei* have diverged with continuous but asymmetric gene flow. Diagram is based on an estimated demographic model for this species pair, with width of boxes proportional to population size and width of arrows is proportional to migration rate (see Table 1).

**Table 1.** Maximum likelihood values and maximum-likelihood parameter estimates for the five demographic models tested.

| Parameters | No<br>migration | Secondary<br>contact - one<br>burst of<br>migration | Continuous<br>migration that<br>starts after<br>divergence | Continuous<br>migration that<br>ends before<br>present day | Continuous<br>migration |
| --- | --- | --- | --- | --- | --- |
| # of parameters estimated | 7 | 7 | 7 | 7 | 6 |
| Ancestral population size | 2075281 | 1962052 | 1971660 | 2006404 | 1982187 |
| *N. pinetum* population size | 645779 | 369032 | 322897 | 337579 | 328311 |
| *N. lecontei* population size | 1057278 | 1052921 | 1104528 | 1075336 | 1093739 |
| Time since divergence (generations) | 1010864 | 1255392 | 1542789 | 1553120 | 1548690 |
| *N. pinetum* bottleneck size | 717 | NA | NA | NA | NA |
| *N. lecontei* bottleneck size | 470 | NA | NA | NA | NA |
| Time since bottle neck | 1010854 | NA | NA | NA | NA |
| Admixture proportion from *N. pinetum* to *N. lecontei* | NA | 0.0060941 | NA | NA | NA |
| Admixture proportion from *N. lecontei* to *N. pinetum* | NA | 0.118292 | NA | NA | NA |
| Time since admixture | NA | 115046 | NA | NA | NA |
| Migration rate from *N. lecontei* to *N. pinetum* | NA | NA | 3.94E-07 | 3.63E-07 | 3.65E-07 |
| Migration rate from *N. pinetum* to *N. lecontei* | NA | NA | 1.64E-08 | 1.83E-08 | 1.71E-08 |
| Number of *N. pinetum* migrants | NA | NA | 0.1273259 | 0.1226254 | 0.1196984 |
| Number of *N. lecontei* migrants | NA | NA | 0.0181423 | 0.0196645 | 0.018665 |
| Time since migration started | NA | NA | 901891 | NA | NA |
| Time since migration ended | NA | NA | NA | 1906 | NA |
| Estimated maximum likelihood (log 10) | -33034.2 | -32805.5 | -32794.6 | -32795 | -32794.3 |
| Maximum observed likelihood (log 10) | -32727.9 | -32727.9 | -32727.9 | -32727.9 | -32727.9 |
| **Maximum Likelihood (est-obs)** | **-306.3** | **-77.6** | **-66.7** | **-67.1** | **-66.4** |

Comparing three different models that included migration (staring after divergence, stopping before the present day, or continuous migration), our SFS data were most likely under the model that had the fewest parameters: a continuous migration model (Table 1). Maximum likelihood parameter estimates under this model suggest that *N. pinetum* and *N. lecontei* diverged ∼1.5×10^6^ generations ago. Assuming 1-3 generations per year for KY populations of these species (Benjamin 1955; Rauf and Benjamin 1980; CL, personal observation), this estimate suggests that *N. pinetum* and *N. lecontei* likely diverged between 0.5 and 1.5 million years ago. Our parameter estimates also suggest that *N. pinetum* has a smaller *N_e_* than *N. lecontei*, and that migration rates have been asymmetric, with more migration from *N. lecontei* to *N. pinetum* than the reverse (Figure 6C, Table1). Importantly, this model provides a good fit to the observed SFS and other summary statistics (Supplementary Figure S6).

Despite continuous migration throughout divergence, genome-wide average F_ST_ was high (F_ST_ = 0.63). However, differentiation levels varied widely across the genome, with localized regions of both very high and very low F_ST_ (Figure 7). Using simulations according to the inferred demographic history to generate 95% confidence intervals for F_ST_ under neutrality revealed evidence of both high-F_ST_ and low-F_ST_ outliers in our empirical dataset (Figure 7). These regions are candidates for divergent selection and adaptive introgression, respectively. Nucleotide diversity (π) for both *N. pinetum* and *N. lecontei* also varied across the genome, but *lecontei* had a higher average π (2.71×10^-5^) than *N. pinetum* (1.95×10^-5^) which is consistent with the differences in effective population size between the two species (Table 1). Overall, demographic modelling and genomic differentiation patterns are consistent with the hypothesis that this species pair diverged with substantial gene flow, while experiencing divergent selection at many unlinked locations throughout the genome.

**Figure 7.**
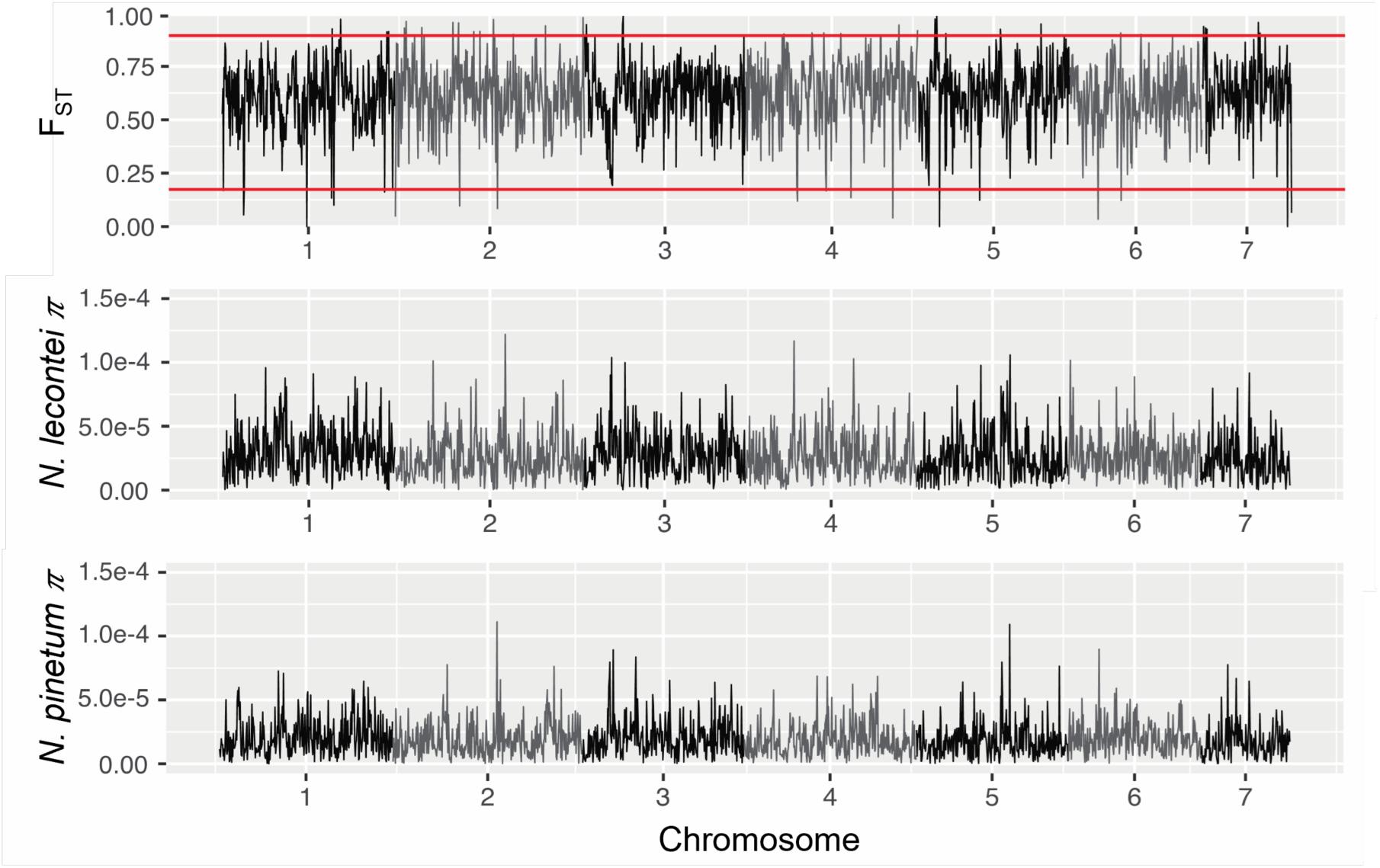
Genome scans of differentiation and diversity for *N. pinetum* and *N. lecontei.* Genetic differentiation (F_ST_) and genetic diversity (nucleotide diversity π) for *N. lecontei* and *N. pinetum* calculated in 100-kb windows. The red lines mark the 95% confidence interval obtained from data simulated under neutrality and the demographic model estimated for this species pair.

### Expectations for faster-haplodiploid effects under inferred demographic model

There were several differences between our simulations and our empirical system, including lower migration rates and asymmetries in both effective population size and migration rate (Table 1) as well as female-biased sex ratios (Harper et al., 2016). To capture some of these system-specific characteristics, we simulated haplodiploid and diploid populations evolving under the demographic model we estimated from our sawfly data. A comparison between our observed summary statistics (SFS, F_ST_, Π, D_xy_, and r^2^) for the putatively neutral intergenic regions and summary statistics obtained from neutral simulations (2*Ns* = 0) shows that the demographic model implemented in SLiM is working as expected, and that our simulated diploid and haplodiploid chromosomes do not differ under neutrality (Supplementary Figure S6).

When we included divergent natural selection in our sawfly-parameterized simulations of diploid and haplodiploid genomes, we once again observed faster-haplodiploid effects on the “genome-wide” mean F_ST_ for both dominant and recessive alleles, rare and common genetic variants, and under a range of selection coefficients (Figure 8). As observed for simulations under the simpler isolation-with-migration model, the magnitude of this effect was highest for rare, recessive alleles and moderate-to-strong selection. These simulations also demonstrate faster-haplodiploid effects can be observed with female-biased sex ratios and with asymmetric migration.

**Figure 8.**
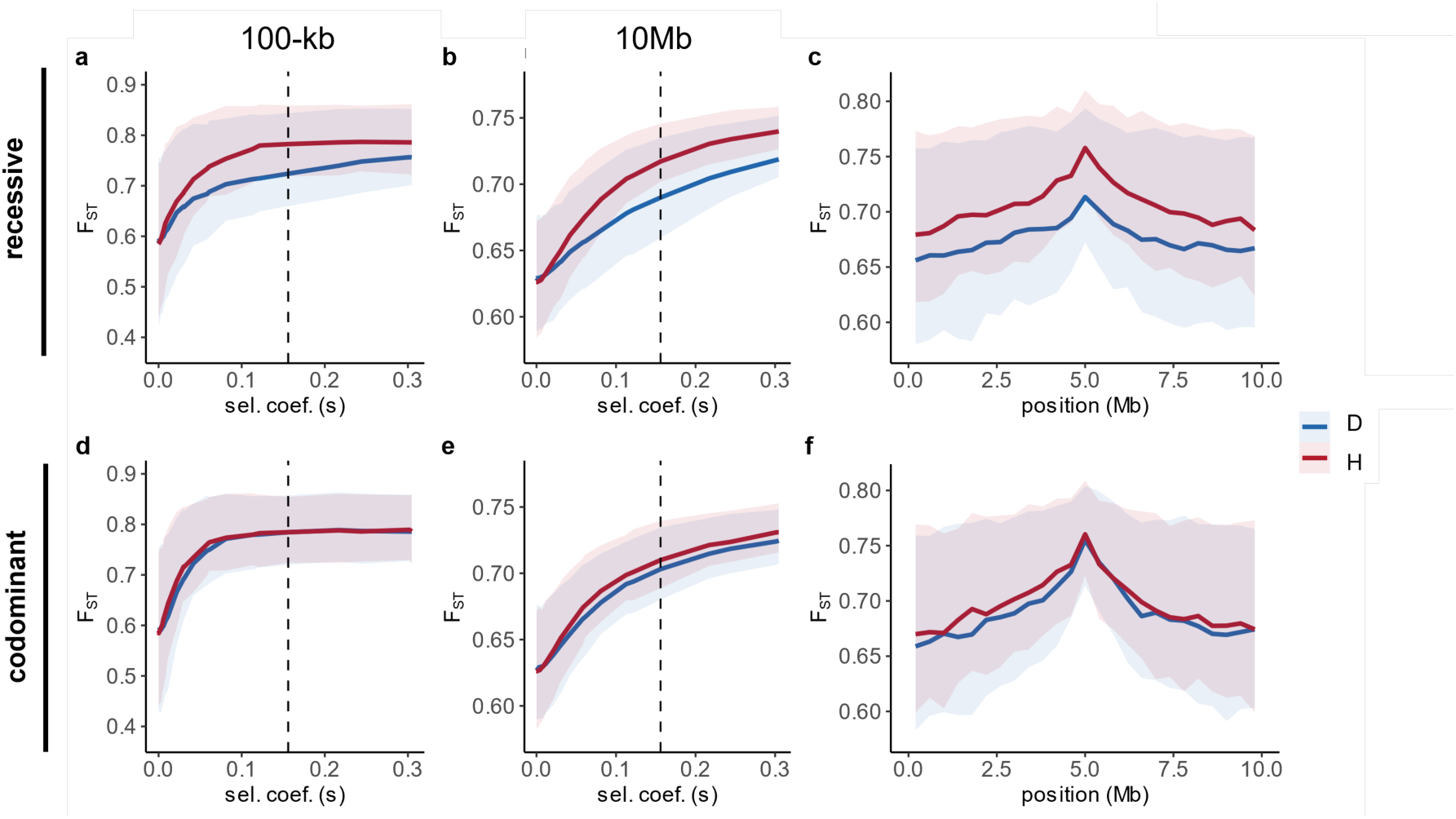
Effect of haplodiploidy and divergent selection on differentiation under inferred demographic history of *Neodiprion* sawflies. Results from simulations performed assuming a sex-ratio with a proportion of 0.7 females and 0.3 males. **(a-b, d-e)** Mean F_ST_ and interquantile range for diploid and haplodiploid populations as a function of selective coefficient for recessive **(a-b)** and codominant **(d-e)** mutations, for different window sizes centered at the selected site **(a,d)** 100Kb, **(b,e)** 10Mb. **(c, f)** Genome scan of F_ST_ for diploid and haplodiploid for recessive **(c)** and codominant **(f)** mutations. Results for *s* = 0.16 with initial frequency of 0.1. Solid line corresponds to mean F_ST_ and shaded area indicates interquantile 0.25-0.75 range. Dashed lines in (**a-b**, **d-e**) indicate the selective coefficient used in genome scan shown in **c, f**.

## Discussion

Haplodiploid taxa are numerous and ecologically diverse (Forbes et al., 2018; Hölldobler & Wilson, 1990). While haplodiploid diversity could be due low transition rates between haplodiploidy and diploidy, it is also possible that haplodiploidy increases speciation rates (Blackmon et al., 2017; Koevoets & Beukeboom, 2008; Lohse & Ross, 2015; Patten et al., 2015). Here we explore one avenue through which haplodiploidy may facilitate speciation: by increasing genomic differentiation and linkage disequilibrium between populations that diverge with gene flow. Specifically, our simulations reveal that so long as there is both selection and migration, haplodiploid populations will maintain higher levels of differentiation than comparable diploid populations. This is true not only at the selected site, but also up to ∼20 cM away. With the sawfly empirical data, we identify a potential case of sympatric divergence via adaptation to different hosts. Here, we discuss implications of these results for faster-X theory, evolution in haplodiploids, and models of sympatric speciation. We also discuss some limitations of our models and data and highlight priorities for future work.

### Faster-haplodiploid effects under divergence-with-gene-flow and relevance to faster-X theory

Overall, our simulations demonstrate multiple mechanisms through which genomic differentiation in haplodiploids is increased relative to diploids when populations diverge with gene flow. Given similarities between the transmission genetics of haplodiploid genomes and X chromosomes, these findings are also relevant to faster-X theory. As described below, our simulations recapitulate several key results from previous work on faster-X theory, albeit in some additional corners of parameter space (e.g., divergence-with-gene-flow via new or rare mutations; *c.f.* Lasne et al., 2017). Based on our simulations, we can group faster-haplodiploid effects and mechanisms into three distinct phases. In the first phase, the dynamics of a divergently selected, low-frequency allele is mostly determined by the risk of loss due to drift (Figure 3, Supplementary Figure S3). The increased efficacy of selection and the reduced probability of allele loss under haplodiploidy and a divergence-with-gene-flow scenario is analogous to classical faster-X divergence among isolated populations in which dominance has a large impact on the outcomes (e.g., Charlesworth et al., 1987; Vicoso and Charlesworth 2009; Meisel and Connallon 2013; Charlesworth et al., 2018). However, assuming sufficient recombination, differential allele loss during phase 1 has a minimal impact on differentiation at linked neutral sites (e.g., compare Figures 2d and 2h to Figures 4e and 5e, respectively).

Once populations escape phase 1 without losing the divergently selected allele, there is a second transitional phase during which the increased efficacy of selection against locally maladaptive alleles reduces effective migration rates and causes haplodiploid loci to differentiate more rapidly than comparable diploid loci (Figures 4 and 5). Again, this is in line with classical faster-X theory demonstrating shorter sojourn times for beneficial X-linked alleles in isolated populations (Avery, 1984; Betancourt et al., 2004). Reduced sojourn times in phase 2 also reduce opportunities for recombination between locally adaptive and maladaptive haplotypes, thereby affecting linked variation (Figures 4 and 5), analogous to predictions for X-linked variation in isolated populations (Betancourt et al., 2004; Owen, 1988).

Once diverging populations approach equilibrium between selection, migration, and drift, they enter phase 3. In this phase, haplodiploidy increases the efficacy of selection against locally maladapted immigrant alleles, resulting in higher allele frequency differences at hemizygous loci compared to diploid loci (Figures 4 and 5). Consistent with deterministic results obtained under similar demographic models (Lasne et al., 2017), we find that when populations remain connected by gene flow, faster-haplodiploid effects occur irrespective of dominance. These findings contrast with classical faster-X theory that has been developed for divergence in isolation, which predicts increased substitution rate only when beneficial mutations are recessive (e.g., Charlesworth et al., 1987; Vicoso & Charlesworth 2006; Meisel & Connallon 2013; Charlesworth et al., 2018) or when codominance is accompanied by deviations from 50:50 sex ratio (Vicoso & Charlesworth, 2009). Additionally, efficient selection against maladapted migrant alleles in Phase 3 causes reduced opportunities for recombination in haplodiploids. This mechanism produces faster-haplodiploid effects at neutral sites linked to both recessive and codominant alleles (Figures 4 and 5). These results are also consistent with predictions from deterministic continent-island models of secondary-contact (Fraïsse & Sachdeva, 2021; Fusco & Uyenoyama, 2011; Muirhead & Presgraves, 2016). Despite several important differences between our model and these secondary contact models, including divergence scenario, migration direction, and the presence of drift, we reach qualitatively similar conclusions. These similarities suggest that in the long term, after migration-selection-drift equilibrium is reached, the impact of the initial phases is negligible.

Where our work departs most from previous faster-X theory is that—to better connect theory to data—we have explicitly modelled the effects of sex-limited hemizygosity and recombination on population genomic datasets. Here, a couple of surprises have emerged. First, under some parameter combinations, faster-haplodiploid effects can be observed in loosely linked neutral sites without corresponding effects at the selected site and tightly linked sites (Figure 5h). These patterns, which are dependent on recombination rate, emerge when haplodiploid populations diverge more rapidly than diploid populations, but ultimately reach the same equilibrium allele frequency. In essence, more rapid differentiation and reduced opportunities for recombination between divergently selected haplotypes in haplodiploids locks linked variation into place, and moderate to strong selection prevents erosion of linkage even with migration. In other words, haplodiploidy leads to larger genomic regions around the selected site with reduced effective migration rate. One important implication of this finding is that haplodiploidy can facilitate the establishment of beneficial mutations that appear in such genomic regions (Yeaman et al., 2016, see below).

Second, while previous work demonstrates that hemizygous selection will give rise to faster-X effects at selected and linked sites, there has been some uncertainty as to how much of the genome is likely to be impacted when there is recurrent migration (Presgraves, 2018). Here, we show that with strong selection (2*Ns*>100) and high migration (2*Nm*∼5), regions of elevated differentiation in haplodiploid chromosomes will be higher and wider than in corresponding diploid chromosomes. Moreover, divergent selection at a single haplodiploid locus can reduce gene flow relative to the diploid case even at neutral sites more than 250-kb away, corresponding to >4.40 cM in simulations under the symmetric isolation-with-migration model (Figures 4 and 5), or >21.5 cM in simulations with sawfly-specific parameters (Figure 8). Additionally, the continent-island models of Fusco & Uyenoyama (2011) and Muirhead & Presgraves (2016) predict that selection at X-linked (and, analogously, haplodiploid) sites can also impact unlinked neutral markers. Assuming that at equilibrium, adaptive divergence-with-gene-flow dynamics can be reasonably well approximated by the continent-island model, we speculate that localized reductions in gene flow surrounding hemizygous loci could extend to the chromosome-wide level.

Although our work better connects theory to empirical data, our models make several simplifying assumptions that could impact patterns of faster-haplodiploid differentiation. Relaxing these assumptions and creating more complex, but realistic, models are therefore potentially fruitful avenues for future research. First, the genetic architecture of adaptation to novel niches is likely much more complex than the simple single-locus model considered here (i.e., adaptation is likely due to many loci with variable effect sizes, dominance coefficients, and non-additive interactions). Although there has been some work on how divergent selection on multiple, possibly interacting, loci impact genomic differentiation (e.g., Yeaman et al., 2016; Aeschbacher et al., 2017), this has not been investigated in the context of hemizygosity (but see Fusco & Uyenoyama 2011; Fraïsse & Sachdeva 2021). Second, the divergence model we considered was also relatively simple, ignoring sex-specific effects such as sex-biased migration, sex-specific selection, and the absence of dosage compensation. For instance, Lasne et al. (2017) predicted that sex-specific migration has a large impact on faster X effects in models with strong migration (*m*>>*s*). Third, we have considered a parallel dominance fitness landscape and the magnitude of faster-X differentiation might differ for other models (but see Lasne et al., 2017). Fourth, we assumed that the fitness of haploid males was equivalent to that of diploid homozygotes. Whether or not this is the case depends on mechanisms of dosage compensation and allelic effects in haploid males, which are not well understood (Gardner 2012; Hitchcock 2021; but see Aron et al., 2005; Dearden et al., 2006; Glastad et al., 2014). Finally, we have ignored the effects that removing deleterious mutations with similar effects across populations may have on patterns of differentiation in haplodiploids and diploids (Charlesworth et al., 1993; Charlesworth et al., 1997). Making precise predictions about chromosome-wide levels in population genomic datasets will require modeling these more complex scenarios, as well as considering local variation in mutation and recombination rate.

### Implications for speciation in haplodiploids

One of the longest running debates in evolutionary biology is over the plausibility and prevalence of sympatric speciation, the evolution of reproductive isolation in absence of geographical isolation (Berlocher & Feder, 2002; Bolnick & Fitzpatrick, 2007; Foote, 2018; Via, 2001). Because of their pronounced host specialization and lifelong association with their host plants, *Neodiprion* sawflies have been hypothesized to undergo sympatric speciation (Bush, 1975a, 1975b; Knerer & Atwood, 1973; Linnen & Farrell, 2010). Although gene flow has been ubiquitous throughout *Neodiprion* divergence (Linnen & Farrell, 2007) and *N. pinetum*’s range is nested within *N. lecontei*’s range, species ranges have changed too much to reconstruct the geographic context of speciation from present day range overlap (Linnen & Farrell, 2010). Here, demographic modeling revealed that the model that best explains patterns of genomic variation in *N. lecontei* and *N. pinetum* does not include a period of isolation (Table 1; Figure 6C). However, distinguishing between models of sympatric divergence and secondary contact is difficult (Sousa & Hey, 2013). This difficulty appears to be true for *N. lecontei* and *N. pinetum* as well: models that included continuous migration either starting after a brief period of isolation (∼64000 generations) or ending right before the present day (∼2000 generations) explained the data nearly as well as a continuous migration model (Table 1). Despite this uncertainty, our top three models and ML parameter estimates all point to a scenario in which gene flow was present throughout all or most of the divergence history of these two species.

Previous work also demonstrates that differences in the pines that *N. lecontei* and *N. pinetum* use are likely to generate divergent selection on many different types of traits, including female oviposition traits, correlated male traits, and larval physiology (Bendall et al., 2020, 2017; Benjamin, 1955; Coppel & Benjamin, 1965; Rauf & Benjamin, 1980; Wilson et al., 1992). Consistent with a “multifarous” or “multidimensional” model of divergent selection (Feder & Nosil, 2010; Rice & Hostert, 1993; White & Butlin, 2021), multiple unlinked loci exceeded expected levels of differentiation under neutrality (Figure 7). We also observed multiple regions of unusually low differentiation, which could be explained by adaptive introgression. We acknowledge, however, that other mechanisms besides divergent selection and adaptive introgression can cause dips and valleys in genome scans (Cruickshank & Hahn, 2014; Ravinet et al., 2017). Thus, interpretation of these genome scans would be improved by characterizing the genomic landscape of recombination and gene density, as well as mapping loci underlying divergently selected traits.

Together with previous work characterizing reproductive barriers in this species pair (Bendall et al., 2017), our demographic modeling and genome scan results support a scenario in which adaptation to different pine trees drove the evolution of reproductive isolation in the presence of substantial gene flow. There is little debate that, given sufficiently strong selection, genetic and phenotypic differences, such as divergently selected host-use traits, can be maintained in the face of gene flow. Instead, the primary objection to sympatric speciation has been that gene flow and recombination will tend to break up associations among favorable combinations of alleles and between divergently selected loci and loci that confer reproductive isolation (Felsenstein, 1981). Multiple mechanisms can help overcome this “selection-recombination” antagonism, thereby aiding the evolution of reproductive isolation when there is gene flow, including pleiotropy (“magic traits” wherein loci underlying local adaptation also confer reproductive isolation (Servedio et al., 2011)) and genomic features that reduce recombination (e.g., chromosomes inversions (Kirkpatrick & Barton, 2006; Ravinet et al., 2017)). Because *Neodiprion* mate on the host plant, it is possible that alleles underlying divergent host preferences also produce habitat isolation (Linnen & Farrell, 2010). However, there is little evidence of chromosomal rearrangements that would reduce recombination in *N. lecontei-N. pinetum* hybrids (Geib and Sim, personal communication).

Even in the absence of “magic traits” and inversions, divergent selection can also facilitate the evolution of reproductive isolation through effects on linked variation (divergence hitchhiking; Via, 2009; Via & West, 2008) and, when there are multiple divergently selected loci, via a genome-wide reduction in the effective migration rate (genome hitchhiking; Barton & Bengtsson, 1986; Feder et al., 2012a,b; Flaxman et al., 2012, 2013). By simulating genomic differentiation under divergent selection, our estimated demographic model, and other system-specific details (sex ratio, recombination rate), we show that haplodiploid inheritance in *N. lecontei* and *N. pinetum* likely increased differentiation at selected and linked loci relative to a comparable diploid scenario (Figure 8). The impact of haplodiploidy was most pronounced for recessive mutations and at intermediate selection coefficients, with effects extending over sizable regions of the genome (comparable to ∼10Mb). By increasing differentiation at linked sites, haplodiploidy could facilitate both divergence hitchhiking and genome hitchhiking, thereby promoting speciation-with-gene-flow. This hypothesis could be tested more directly via simulations that examine the impact of haplodiploidy on non-neutral linked variation and interactions between multiple loci (Feder et al., 2012b; Feder & Nosil, 2010; Flaxman et al., 2012; Nosil & Feder, 2012; Via, 2012; Yeaman & Whitlock, 2011; Yeaman et al., 2016).

### Conclusions

Overall, our work suggests that sex-limited hemizygosity and recombination, both of which are maximized in Hymenoptera and other haplodiploid clades, can have substantial effects on genomic differentiation in wild populations. One potential implication of this work is that haplodiploid taxa are more likely to undergo sympatric speciation and can withstand greater levels of gene flow during divergence, which may ultimately give rise to higher rates of local adaptation and speciation. A comparative analysis of divergence history between diploids and haplodiploids would be an informative step in testing this hypothesis. More generally, there are potentially numerous evolutionary consequences of haplodiploidy that may shed light on patterns of biodiversity and have implications that extend to non-haplodiploid taxa.

## Acknowledgements

We thank members of the Linnen lab for assistance with collection and rearing of pine sawflies, and we thank Kim Vertacnik for providing data from an outgroup taxon. This work was supported by the National Science Foundation (DEB-CAREER-1750946 and DEB-1257739 to C.R.L); USDA-NIFA pre-doctoral fellowship (2015-67011-22803 to R.K.B.); Portuguese National Science Foundation (“Fundação para a Ciência e a Tecnologia” – FCT; UIDB/00329/2020, individual grants CEECIND/02391/2017 and CEECINST/00032/2018/CP1523/CT0008 to V.C.S.); and by EU H2020 program (Marie Skłodowska-Curie grant number 799729 to V.C.S.). For computing resources, we thank the University of Kentucky Center for Computational Sciences and the Lipscomb High Performance Computing Cluster, as well as the INCD (https://incd.pt/), for which access was funded through FCT Advanced Computing Projects (CPCA/A0/7303/2020 to V.C.S.).

## Data accessibility statement

All the scripts used for SLiM v3 simulations (bash and Slim input files) and analysis of SLiM results (R scripts) are available on VCS’s GitHub page (https://github.com/vsousa/EG_cE3c) ([dataset] Bendall et al., 2021a). Upon acceptance for publication, *Neodiprion* sequencing reads will be deposited in the NCBI SRA ([dataset] Bendall et al., 2021b). All VCF files, custom scripts, and input files for the analysis of sawfly data will be deposited in DRYAD ([dataset] Bendall et al., 2021c).

## Author contributions

EEB, VCS, and CRL designed the research; EEB, VCS, RKB, and CRL performed the research; EEB, VCS, and CRL analyzed the data; EEB, VCS, and CRL wrote the paper, with input from all authors.

