## Supplementary Text, Table and Figures for "Faster-haplodiploid evolution under divergence-with-gene-flow: simulations and empirical data from pine-feeding hymenopterans"

### Supplemental Methods

#### Scaling of haplodiploid and diploid SLIM simulations

In a population with $N_F$ females and $N_M$ males, the effective population size of diploid autosomes is $N_D$=(4$N_M N_F$)/($N_M$+$N_F$)) and of haplodiploid chromosomes is $N_H$=(9$N_M N_F$)/(4$N_M$+2$N_F$) (Wright 1969, Mendez 2017). Thus, the ratio $x=N_H/N_D$ of haplodiploid to diploid effective sizes depends on the number of individuals from each sex. Defining the sex-ratio as the proportion of males $sr=N_M/(N_M+N_F)$, $x=N_H/N_D$ simplifies to $x$=(9/8)(1/(1+$sr$)), recovering $x$=3/4 for a sex-ratio of 50:50 ($sr$=0.50). We adjusted the $N_e$ to ensure that both the diploid and haplodiploid chromosomes have 2$N_e$=1500. This is the effective size of a hemizygous locus with $N$=1000 individuals (500 females with two copies and 500 males with one copy). This corresponds in the diploid case to $N_D=x\,N$ individuals, where $x$=3/4 for a 0.50 sex-ratio, resulting in $N_D$=750 individuals ($N_F$=375 females and $N_M$=375 males). To ensure that the average recombination rate was the same for diploid and haplodiploid chromosomes, we multiplied the diploid recombination rate by 2/3 since the haplodiploid chromosome spends 2/3 of its time in the sex in which it recombines (Lohmueller et al., 2010). This scaling is important because average recombination rate can impact linked variation and there is evidence that haplodiploid genomes have average recombination rates comparable to autosomes, despite the presence of non-recombining males (Kong et al., 2002; Wilfert et al., 2007).

#### Scaling of SLIM simulations for *Neodiprion* sawflies

To obtain the number of individuals $N$ in SLiM that correspond to the haploid effective sizes estimated with fastsimcoal2, while accounting for a sex-ratio $sr=N_M/(N_F+N_M)$=0.30 we used the expressions $N_e$=(4$N_M N_F$)/($N_M$+$N_F$)) for diploids and $N_e$=(9$N_M N_F$)/(4$N_M$+2$N_F$) for haplodiploids (Wright 1969; Mendez 2017), where $N_F$ is the number of females and $N_M$ is the number of males. Noting that $N=N_F+N_M$ and replacing $N_M=srN$ and $N_F=(1-sr)N$ these expressions simplify to $N_e=4Nsr(1-sr)$ and $N_e=(9Nsr(1-sr))/(2(1+sr))$. Solving for the number of individuals we obtain $N=N_e/(4sr(1-sr))$ for diploids and $N=(2(1+sr)N_e)/(9sr(1-sr))$ for haplodiploids.

### Supplemental Figures

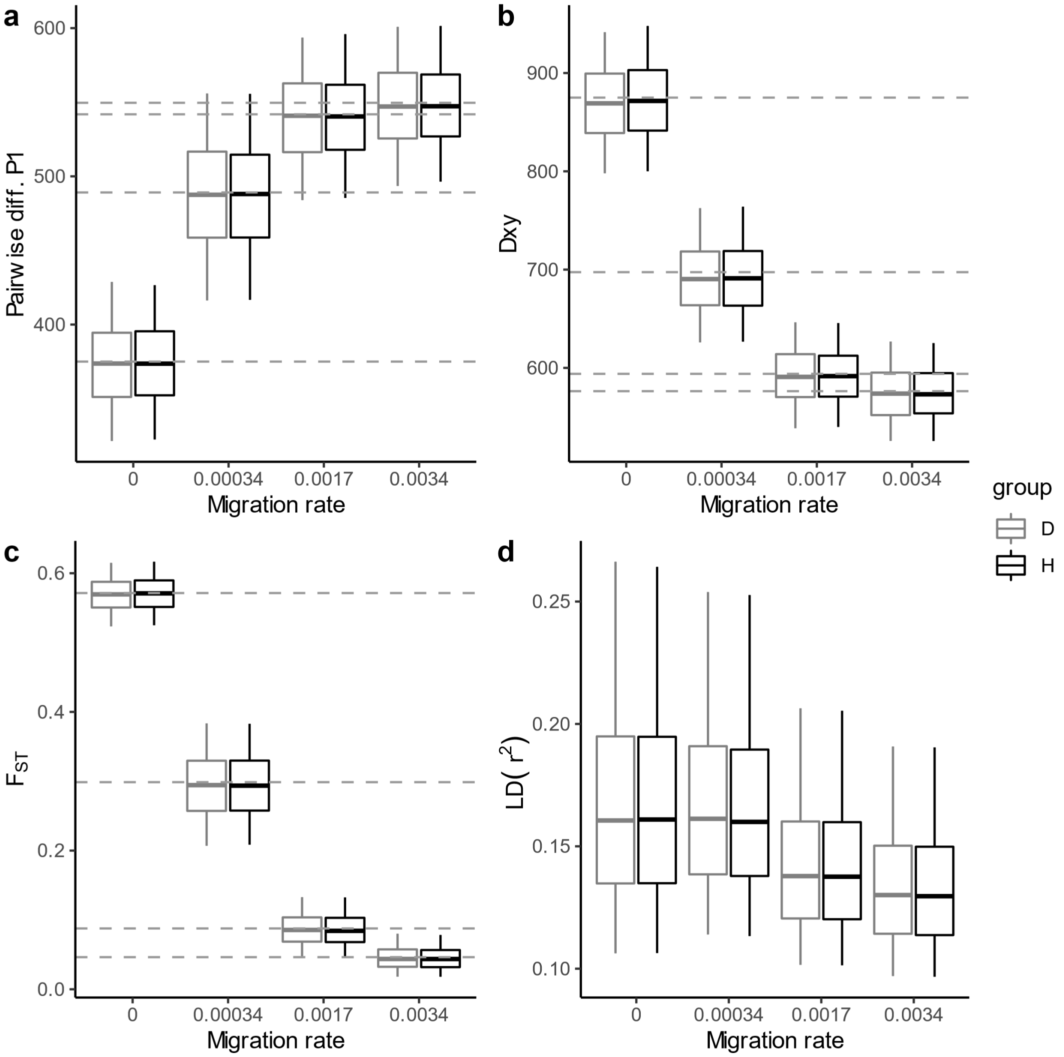


**Supplementary Figure S1. Comparison of diploid and haplodiploid scaled neutral (*s*=0) SLiM simulations to each other and to theoretical expectations**. Each plot shows the distribution of summary statistics obtained under neutrality (*s*=0) for different migration rates, pooling simulations with initial frequencies of $q_0$=(1/(2$N_e$), 0.01, 0.1, 0.5), comparing diploid (D) and haplodiploid (H) cases with 500-kb chromosomes. Whiskers of boxplots extend to 0.05 and 0.95 qua ntiles. Dashed horizontal lines indicate expected values under neutrality obtained for an isolation-with-migration model based on within- and between-population coalescent times (eq. 22 of Wilkinson-Herbots 2008). **A)** Nucleotide diversity (pi for population 1). **B)** Dxy. **C)** $F_{ST}$. **D)** Linkage disequilibrium measured with mean $r^2$ across all pairs of SNPs in windows of 20Kb. The scaled simulations with SLiM fit the neutral expectations, and there are no differences in diversity, differentiation and linkage disequilibrium patterns between diploid and haplodiploid simulations. Furthermore, this shows that the parameter combinations cover a wide range of diversity and differentiation values.

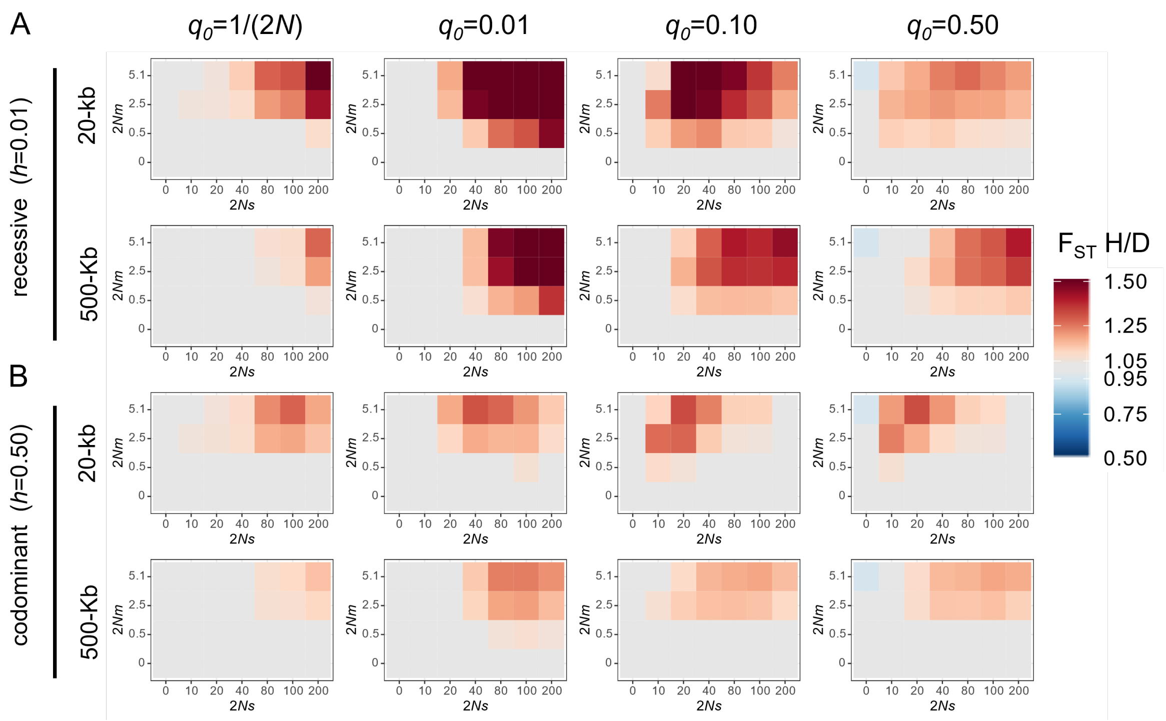


**Supplementary Figure S2. Faster-haplodiploid effects as a function of initial frequency of derived allele.** Heatmaps of the ratio of average haplodiploid to diploid (H/D) differentiation ($F_{ST}$) with different initial frequencies $q_0$ for **A**) recessive (*h*=0.01) alleles; and **B**) codominant *(h*=0.50) alleles*.* Results were obtained for the average $F_{ST}$ across 1000 simulations, based on a sample of 20 gene copies from each population. The heatmap $F_{ST}$ H/D scale was truncated: values larger than 1.5 were assigned the darkest red color, and all values lower than 0.5 were assigned the same blue color.

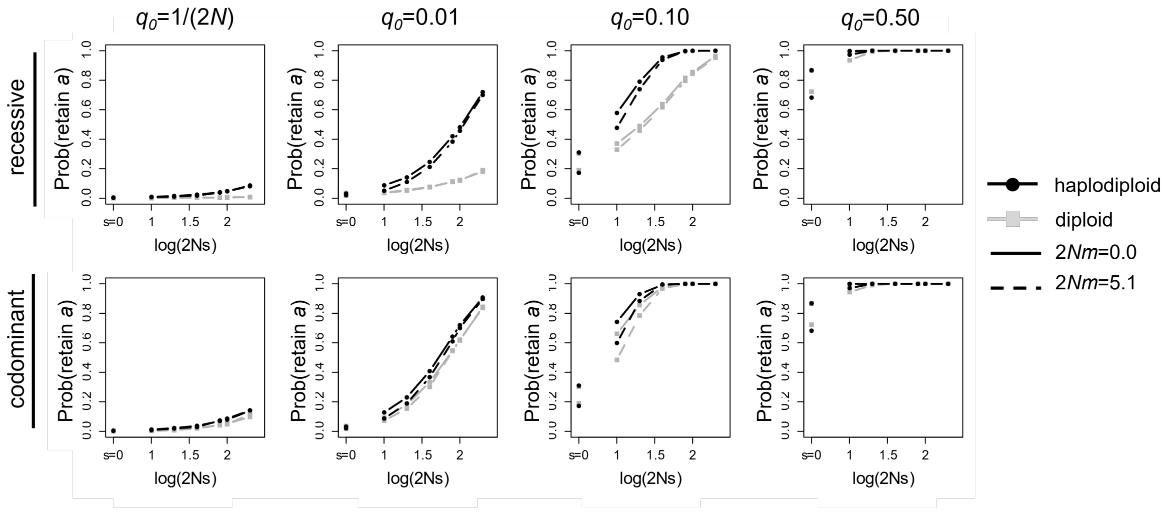


**Supplementary Figure S3. Effect of haplodiploidy on the probability of retaining the derived allele.** Probability of retaining the derived allele *a* for haplodiploid and diploid populations at the two extremes of migration rates considered: no migration (2*Nm*=0, solid line) and high migration (2*Nm*=5.1, dotted line). Each column corresponds to a different initial allele frequency ($q_0$) and rows correspond to recessive and codominant mutations, respectively.

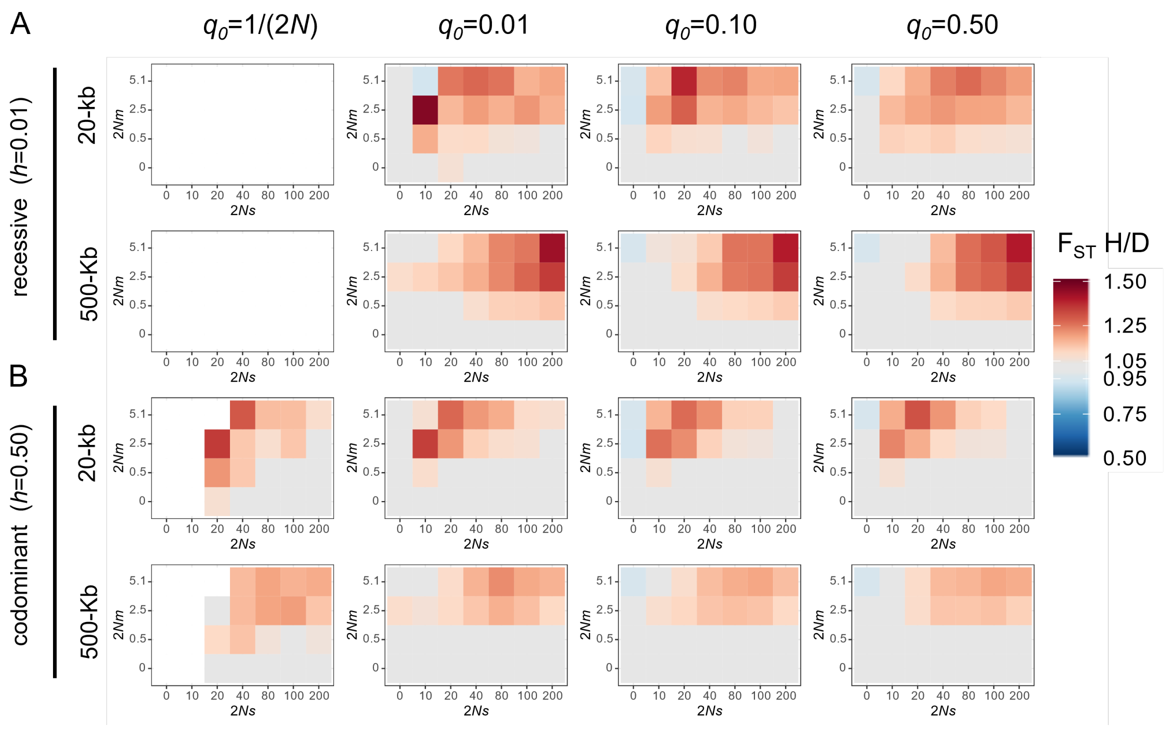


**Supplementary Figure S4. Faster-haplodiploid effects conditional on retaining the derived selected allele under different initial allele frequencies.** Heatmaps of ratio of average haplodiploid to diploid (H/D) differentiation ($F_{ST}$) with different initial frequencies $q_0$ for **A**) recessive ($h$=0.01) mutations; and **B**) codominant *(h*=0.50) mutations*.* Results obtained for the average $F_{ST}$ across simulations that kept the derived allele after 2,000 generations, based on a sample of 20 gene copies from each population. Average $F_{ST}$ for cases with less than 10 simulations was treated as missing data (shown in white). The heatmap $F_{ST}$ H/D scale was truncated: values larger than 1.5 were assigned the red color, and all values lower than 0.5 were assigned the same blue color.

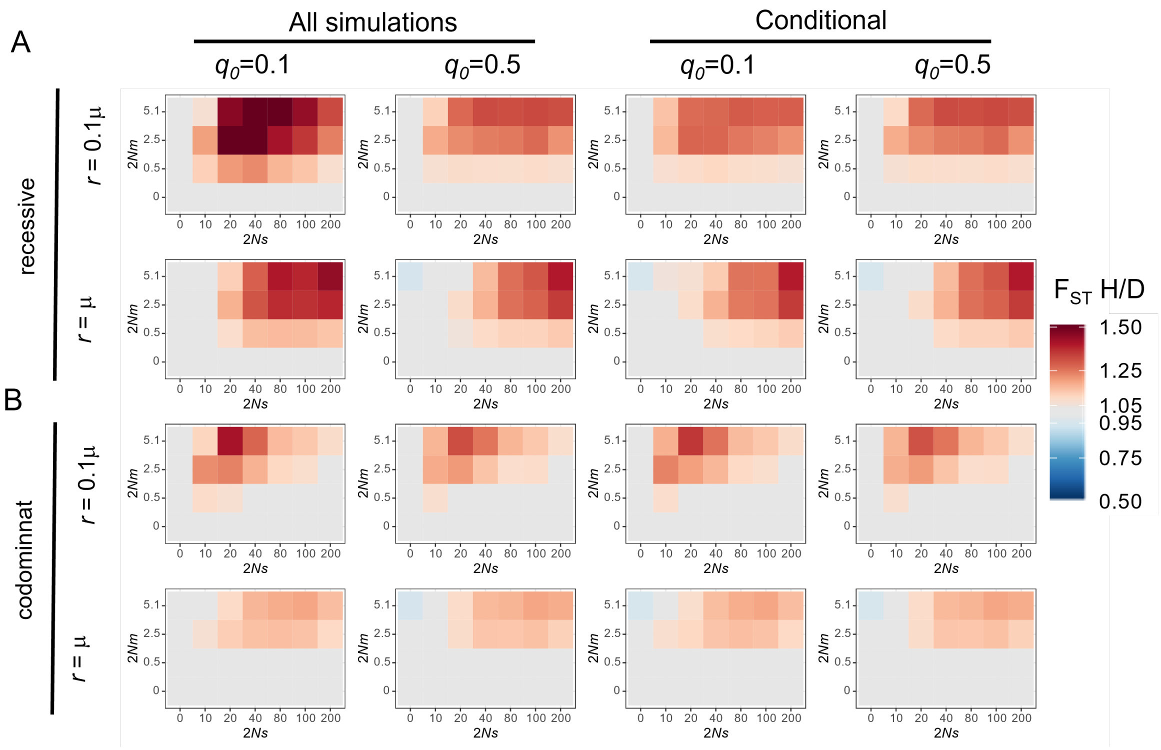


**Supplementary Figure S5. Effect of recombination rate on faster haplodiploid evolution.** Ratio of haplodiploid to diploid (H/D) average $F_{ST}$ for recessive and codominant mutations with different initial frequencies $q_0$ and recombination rates $r$ (in relation to the mutation rate μ). Results of average $F_{ST}$ ratios **A)** across all 1000 simulations for each parameter combination; and **B)** conditional on simulations retaining the derived allele *a* in population 1. The heatmap $F_{ST}$ H/D scale was truncated: values larger than 1.5 were assigned the red color, and all values lower than 0.5 were assigned the same blue color.

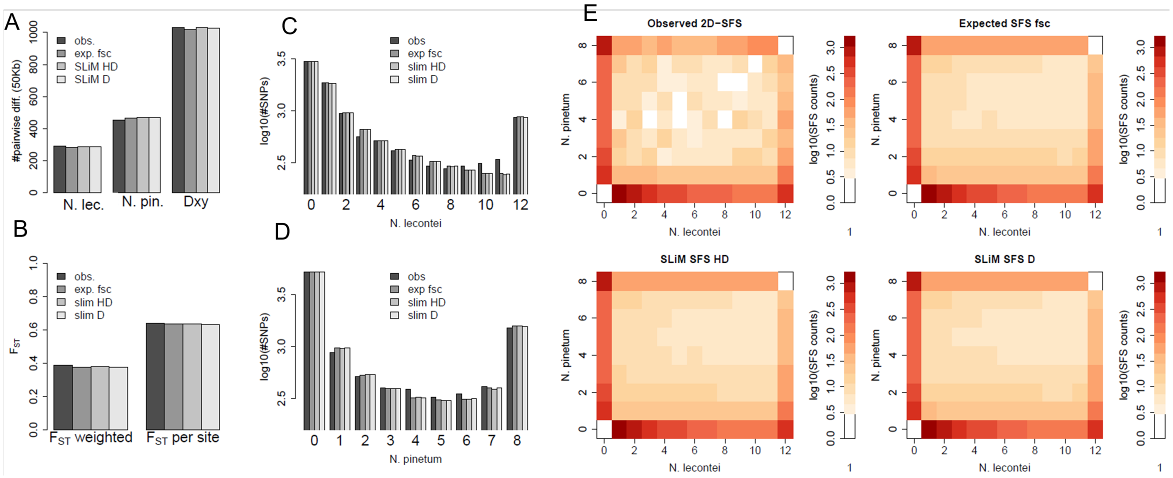


**Supplementary Figure S6. Comparison of neutral simulations done with fastsimcoal2 and SLiM to observed *Neodiprion* sawfly data**. **A)** Fit of within population diversity (number of pairwise differences) and Dxy. **B)** Fit of $F_{ST}$ computed over all SNPs ($F_{ST}$ weighted) or per site ($F_{ST}$ per site). **C)** Fit of marginal 1D-SFS for *N. lecontei*. **D)** Fit of marginal 1D-SFS for *N. pinetum*. **E)** Observed 2D-SFS for *Neodiprion* and 2D-SFS obtained according to the best-fit model with fastsimcoal2 and SLiM. Observed statistics (obs) were compared to simulations done with the parameter estimates obtained with fastsimcoal2 (exp. fsc) for the isolation with migration model with continuous migration, and with SLiM simulations done for diploid (slim D) and haplodiploid (slim HD). For SLiM, the simulations were done re-scaling the parameters such that it was computationally efficient, furthermore the effective sizes and recombination rates were re-scaled such that the neutral diversity and LD patterns were similar between diploids and haplodiploids. Results show that the re-scaling does not affect the summary statistics considered, and that under neutrality, the haplodiploid (slim HD) and diploid (slim D) simulations produce nearly identical summary statistics.

### Supplemental Tables

**Supplementary Table S1**. Combination of parameters used for simulations under the symmetric isolation with migration model.

| | Stochastic simulations (SLiM) | |
|---|---|---|
| | **diploid** | **haplodiploids** |
| **Ancestral population** | 10,000 | 10,000 |
| **Time of split (generations)** | 2,000 | 2,000 |
| **Effective size (2$N_e$)** | 1,500 | 1,500 |
| **Number of individuals (SLiM)** | 750 | 1,000 |
| **Mutation rate ($\mu$)** | 2.5x$10^{-7}$ | 2.5x$10^{-7}$ |
| **Recombination rate (*r*)** | (2/3) 2.5x$10^{-7}$ | (2/3) 2.5x$10^{-7}$* |
| | (2/3) 2.5x$10^{-8}$ | (2/3) 2.5x$10^{-8}$* |
| **Initial frequency $q_0$** | 1/(2$N_e$)<br>0.01<br>0.1<br>0.5 | 1/(2$N_e$)<br>0.01<br>0.1<br>0.5 |
| **Dominance of allele a (h)** | 0.01<br>0.5 | 0.01<br>0.5 |
| **Migration rate (*m*)** | 0 (2*Nm*=0.0)<br>0.00034 (2*Nm*=0.5)<br>0.0017 (2*Nm*=2.5)<br>0.0034 (2*Nm*=5.1) | 0 (2*Nm*=0.0)<br>0.00034 (2*Nm*=0.5)<br>0.0017 (2*Nm*=2.5)<br>0.0034 (2*Nm*=5.1)) |
| **Selection coefficient (*s*)** | 0 (2*Ns*=0)<br>0.00670 (2*Ns*=10)<br>0.01333 (2*Ns*=20)<br>0.02667 (2*Ns*=40)<br>0.05333 (2*Ns*=80)<br>0.06670 (2*Ns*=100)<br>0.13340 (2*Ns*=200) | 0 (2*Ns*=0)<br>0.00670 (2*Ns*=10)<br>0.01333 (2*Ns*=20)<br>0.02667 (2*Ns*=40)<br>0.05333 (2*Ns*=80)<br>0.06670 (2*Ns*=100)<br>0.13340 (2*Ns*=200) |

**Supplementary Table S2.** Sampling locations, population identification, and barcodes for *Neodiprion* sawflies.

| Individual | Species | Population | Latitude | Longitude | Index | Barcode |
|---|---|---|---|---|---|---|
| NP002L_01 | *N. pinetum* | KY | 38.026 | -84.495 | TGACCAAT | GCAAGCCAT |
| NP005L_01 | *N. pinetum* | KY | 38.042 | -84.442 | CCGTCCCG | CGTCGCCACT |
| NP006_08 | *N. pinetum* | KY | 38.042 | -84.442 | GTGAAACG | GCGTCCT |
| NP015_01 | *N. pinetum* | KY | 38.016 | -84.490 | TTAGGCAT | TGGCACAGA |
| NP018_V1 | *N. pinetum* | KY | 38.003 | -84.525 | GTGAAACG | GCAAGCCAT |
| NP019_01 | *N. pinetum* | KY | 38.003 | -84.525 | CCGTCCCG | ATATCGCCA |
| NP021L_01 | *N. pinetum* | KY | 38.857 | -84.618 | TGACCAAT | ATAGAT |
| NP022L_01 | *N. pinetum* | KY | 38.857 | -84.618 | CGATGTAT | CGTCGCCACT |
| NP023L_01 | *N. pinetum* | KY | 38.857 | -84.618 | TGACCAAT | AACTGG |
| NP024L_01 | *N. pinetum* | KY | 38.857 | -84.618 | CCGTCCCG | ACAACCAACT |
| NP025L_01 | *N. pinetum* | KY | 38.857 | -84.618 | CCGTCCCG | TATTCGCAT |
| NP026L_01 | *N. pinetum* | KY | 38.857 | -84.618 | GTGAAACG | GGAACGA |
| NP027L_01 | *N. pinetum* | KY | 38.857 | -84.618 | TTAGGCAT | TAGCCAA |
| NP028L_01 | *N. pinetum* | KY | 38.857 | -84.618 | GTGAAACG | ACGGTACT |
| NP029L_01 | *N. pinetum* | KY | 38.857 | -84.618 | ATCACGAT | TATGT |
| NP030L_01 | *N. pinetum* | KY | 38.857 | -84.618 | GTCCGCAC | CCTCG |
| NP034_02 | *N. pinetum* | KY | 38.032 | -84.565 | ATCACGAT | CGTGGACAGT |
| NP036_04 | *N. pinetum* | KY | 38.032 | -84.565 | TTAGGCAT | TGACGCCA |
| NP037L_01b | *N. pinetum* | KY | 38.032 | -84.565 | GTCCGCAC | GGTGT |
| NP038_01 | *N. pinetum* | KY | 38.003 | -84.525 | TGACCAAT | ACAACCAACT |
| NP040_02 | *N. pinetum* | KY | 37.973 | -84.500 | CGATGTAT | TCACGGAAG |
| NP041_01 | *N. pinetum* | KY | 37.973 | -84.500 | TTAGGCAT | AAGACGCT |
| NP047_01 | *N. pinetum* | KY | 39.008 | -84.650 | GTCCGCAC | TATTCGCAT |
| LL002_01 | *N. lecontei* | KY | 38.014 | -84.504 | GTGAAACG | GAGCGACAT |
| LL006_02b | *N. lecontei* | KY | 38.014 | -84.504 | GTGAAACG | TCAGAGAT |
| LL010 | *N. lecontei* | KY | 36.928 | -84.619 | GTGAAACG | AACTGG |
| LL011 | *N. lecontei* | KY | 37.071 | -84.211 | AGTCAACA | CTAAGCA |
| LL013 | *N. lecontei* | KY | 37.071 | -84.211 | TGACCAAT | ACAACT |
| LL014 | *N. lecontei* | KY | 37.071 | -84.211 | CCGTCCCG | ACTGCGAT |
| LL015_01 | *N. lecontei* | KY | 37.071 | -84.211 | ATCACGAT | TCAGAGAT |
| LL053 | *N. lecontei* | KY | 38.014 | -84.504 | TGACCAAT | TCTTGG |
| LL064 | *N. lecontei* | KY | 38.014 | -84.504 | TTAGGCAT | ACGGTACT |
| LL074_1R | *N. lecontei* | KY | 38.014 | -84.504 | AGTCAACA | TCAGAGAT |
| LL097 | *N. lecontei* | KY | 38.044 | -84.497 | GTGAAACG | GGCTTA |
| LL098_02 | *N. lecontei* | KY | 38.044 | -84.497 | ATCACGAT | AACTGG |
| LL099_02 | *N. lecontei* | KY | 38.044 | -84.497 | TGACCAAT | ATTAT |
| LL100 | *N. lecontei* | KY | 38.023 | -84.494 | GTCCGCAC | TGGCACAGA |

| | | | | | | |
|---|---|---|---|---|---|---|
| LL110_02 | *N. lecontei* | KY | 38.024 | -84.532 | ATCACGAT | TCTTGG |
| LL111_02 | *N. lecontei* | KY | 38.024 | -84.532 | TTAGGCAT | ATTAT |
| LL112 | *N. lecontei* | KY | 38.024 | -84.532 | CCGTCCCG | TAGCCAA |
| LL113 | *N. lecontei* | KY | 38.024 | -84.532 | GTCCGCAC | TGGCAACAGA |
| LL117 | *N. lecontei* | KY | 37.984 | -84.418 | CCGTCCCG | CACCA |
| LL123 | *N. lecontei* | KY | 38.033 | -84.507 | TGACCAAT | CCTCG |
| LL124 | *N. lecontei* | KY | 38.033 | -84.507 | ATCACGAT | ATATCGCCA |
| LL129 | *N. lecontei* | KY | 38.044 | -84.497 | TTAGGCAT | CCGAACA |
| LL130 | *N. lecontei* | KY | 38.044 | -84.497 | AGTCAACA | AACGTGCCT |
| LL133 | *N. lecontei* | KY | 38.024 | -84.532 | CGATGTAT | CTAAGCA |
| LL160 | *N. lecontei* | KY | 38.014 | -84.504 | TTAGGCAT | ATAGAT |
| LL181 | *N. lecontei* | KY | 38.402 | -85.586 | GTGAAACG | CACCA |
| LL194_02 | *N. lecontei* | KY | 37.806 | -83.678 | TGACCAAT | TGCTT |
| LL195_02 | *N. lecontei* | KY | 37.806 | -83.678 | GTCCGCAC | CTTGA |
| RB017.05 | *N. lecontei* | KY | 37.984 | -84.511 | AGTCAACA | TAGCCAA |
| RB020Db | *N. lecontei* | KY | 37.066 | -84.159 | GTCCGCAC | ATTAT |
| RB022C | *N. lecontei* | KY | 38.024 | -84.494 | TTAGGCAT | TCACTG |
| RB040_02 | *N. lecontei* | KY | 38.209 | -84.390 | GTGAAACG | ATGAGCAA |
| RB076_01 | *N. lecontei* | KY | 38.014 | -84.504 | GTGAAACG | TGACGCCA |
| RB129_01 | *N. lecontei* | KY | 38.014 | -84.504 | TTAGGCAT | TCAGAGAT |
| RB144 | *N. lecontei* | KY | 38.209 | -84.390 | AGTCAACA | TCACTG |
| RB148 | *N. lecontei* | KY | 38.209 | -84.390 | TTAGGCAT | GAAGTG |
| RB149_02 | *N. lecontei* | KY | 38.209 | -84.390 | ATCACGAT | TAGCCAA |
| RB151 | *N. lecontei* | KY | 38.209 | -84.390 | GTGAAACG | GCCTACCT |
| RB161_02 | *N. lecontei* | KY | 37.984 | -84.418 | GTCCGCAC | ACTGCGAT |
| RB162 | *N. lecontei* | KY | 38.024 | -84.494 | GTGAAACG | AAGACGCT |
| RB344 | *N. lecontei* | KY | 38.014 | -84.504 | AGTCAACA | CCTTGCCATT |
| RB358 | *N. lecontei* | KY | 38.014 | -84.504 | TTAGGCAT | CTTGA |
| RB361 | *N. lecontei* | KY | 37.984 | -84.418 | AGTCAACA | AACTGG |
| RB370_02 | *N. lecontei* | KY | 38.033 | -84.507 | GTCCGCAC | AACTGG |
| RB099_01 | *N. lecontei* | UPMI | 45.924 | -86.303 | ATCACGAT | CAGATA |
| RB405 | *N. lecontei* | UPMI | 45.949 | -86.261 | GTGAAACG | ACTGCGAT |
| RB406 | *N. lecontei* | UPMI | 45.949 | -86.261 | TTAGGCAT | ACAACCAACT |
| RB095_04 | *N. lecontei* | UPMI | 46.094 | -85.339 | TGACCAAT | TCAGAGAT |
| RB096B | *N. lecontei* | UPMI | 46.096 | -85.394 | GTGAAACG | TAGCCAA |
| RB098_01 | *N. lecontei* | UPMI | 46.096 | -85.394 | CCGTCCCG | AACGTGCCT |
| RB480_02 | *N. lecontei* | UPMI | 46.096 | -85.394 | GTCCGCAC | ACAACCAACT |
| RB480_1 | *N. lecontei* | UPMI | 46.096 | -85.394 | TGACCAAT | CGTGGACAGT |
| RB481 | *N. lecontei* | UPMI | 46.096 | -85.394 | AGTCAACA | CGTCGCCACT |
| RB482 | *N. lecontei* | UPMI | 46.096 | -85.394 | GTGAAACG | TCTTGG |

| | | | | | | |
|---|---|---|---|---|---|---|
| RB483 | *N. lecontei* | UPMI | 46.096 | -85.394 | TGACCAAT | CGTCGCCACT |
| RB407 | *N. lecontei* | UPMI | 45.918 | -86.313 | TTAGGCAT | CCACTCA |
| RB408 | *N. lecontei* | UPMI | 45.918 | -86.313 | CGATGTAT | CGTGGACAGT |
| RB409 | *N. lecontei* | UPMI | 45.926 | -86.294 | TTAGGCAT | ACCAGGA |
| RB410 | *N. lecontei* | UPMI | 45.926 | -86.294 | TGACCAAT | GGCTTA |
| RB411 | *N. lecontei* | UPMI | 45.926 | -86.294 | GTCCGCAC | TAGCCAA |
| RB412 | *N. lecontei* | UPMI | 45.926 | -86.294 | AGTCAACA | CGTGGACAGT |
| RB413 | *N. lecontei* | UPMI | 45.926 | -86.294 | CCGTCCCG | GGTGCACATT |
| LL198 | Wild Hybrid | KY | 38.857 | -84.618 | TTAGGCAT | CGTCGCCACT |
| LL244 | Wild Hybrid | KY | 38.014 | -84.504 | ATCACGAT | GGTGCACATT |
| NP046 | Wild Hybrid | KY | 38.893 | -84.557 | GTGAAACG | CTCTCGCAT |
| S23 F1 | Lab reared hybrid | NA | NA | NA | TGACCAAT | CTAAGCA |

**Supplementary Table S3**. Cross validation (CV) error scores for admixture models of *K*=1-5.

| K | CV error |
|---|---|
| 1 | 0.4853375 |
| 2 | 0.3911959 |
| 3 | 0.4113284 |
| 4 | 0.4259785 |
| 5 | 0.440888 |

**Supplementary Table S4**. Combination of parameters used for simulations of haplodiploids and diploids under divergent selection and the asymmetric continuous migration model inferred for *Neodiprion pinetum* (pop. 1) and *N. lecontei* (pop. 2) sawflies, assuming a sex-ratio of 70 females:30 males. See details in Supplementary Methods.

| | **Inferred based on SFS** | **Re-scaled parameters** | **SLiM haplodiploids** | **SLiM diploids** |
|---|---|---|---|---|
| Ancestral population $N_e$ | 1,982,187 | 1,982 | 1,982 | 1,982 |
| Number of individuals (SLiM) ancestral pop | NA | NA | 1,363 | 1,180 |
| Time of split (generations) | 1,548,690 | 1,549 | 1,549 | 1,549 |
| Effective size ($2N_e$) Pop 1 | 328,311 | 328 | 328 | 328 |
| Number of individuals (SLiM) Pop1 | NA | NA | 226 | 195 |
| Effective size ($2N_e$) Pop 2 | 1,093,739 | 1,094 | 1,094 | 1,094 |
| Number of individuals (SLiM) Pop2 | NA | NA | 752 | 651 |
| Mutation rate per site per generation ($\mu$) | $3.5x10^{-9}$ | $3.5x10^{-7}$ | $3.5x10^{-7}$ | $3.5x10^{-7}$ |
| Recombination rate per site per generation ($r$) | NA | $(2/3)*3.5x10^{-7}$ | $2.34x10^{-7}$ | $2.34x10^{-7}$ |
| | | $(2/3)*1.05x10^{-6}$ | $7.00\ x10^{-7}$ | $7.00\ x10^{-7}$ |
| Initial frequency $q_0$ | 0.1 | 0.1 | 0.1 | 0.1 |
| Dominance of allele a ($h$) | NA | NA | 0.01<br>0.5 | 0.01<br>0.5 |
| Immigration rate from *N. lecontei* into *N. pinetum* ($m_{12}$) | $3,65x10^{-7}$ | $3,65x10^{-4}$ | $3,65x10^{-4}$ | $3,65x10^{-4}$ |
| Immigration rate from *N. pinetum* into *N. lecontei* ($m_{21}$) | $1,71x10^{-8}$ | $1,71x10^{-5}$ | $1,71x10^{-5}$ | $1,71x10^{-5}$ |
| Selection coefficient ($s$) | 0 | 0 | 0-0.3 | 0-0.3 |
| Number of sites | 225,277 callable sites (invariant+ SNPs) | 500,000 (corresponding to 50,000 callable sites) | 500,000 (corresponding to 10Mb of ddRAD data) | 500,000 (corresponding to 10Mb of ddRAD data) |
